# Comparative transcriptome analysis of noble crayfish and marbled crayfish immune response to *Aphanomyces astaci* challenges

**DOI:** 10.1101/2021.05.25.445163

**Authors:** Ljudevit Luka Boštjančić, Caterina Francesconi, Christelle Rutz, Lucien Hoffbeck, Laetitia Poidevin, Arnaud Kress, Japo Jussila, Jenny Makkonen, Barbara Feldmeyer, Miklós Bálint, Odile Lecompte, Kathrin Theissinger

## Abstract

Introduction of invasive North American crayfish species and their pathogen *Aphanomyces astaci* has significantly contributed to the decline of European freshwater crayfish populations. In this study, noble crayfish, a susceptible native European species, and marbled crayfish, an invasive disease-resistant species, were challenged with haplogroup A (low virulence) and haplogroup B (high virulence) strain of *A. astaci*. Hepatopancreatic tissue was isolated 3 and 21 days post-challenge. Our results revealed strong up-regulation in expression levels of the prophenoloxidase cascade immune-related genes in the haplogroup B challenged noble crayfish 3 days post-challenge. In the marbled crayfish, we observed an up-regulation of immune system relevant genes (DSCAM, AP, ALFs, CTLs and hemocyanin) 3 days post-challenge. This response highlights the marbled crayfish capability of building the immune tolerance. Furthermore, we successfully characterised several novel immune related gene groups in both crayfish species, contributing to our current understanding of crayfish immune related genes landscape.

**Graphical abstract:** **a)** Study species noble crayfish (*Astacus astacus*) in purple and marbled crayfish (*Procambarus virginalis*) in green challenged with the pathogen *Aphanomyces astaci* haplogroup A (Hap A) strain of low virulence and haplogroup B (Hap B) strain of high virulence. **b)** Sampling scheme of the infection experiment: 5 individuals were taken from the experiment three- and 21-days post-challenge. From each individual, a hepatopancreas sample was taken, followed by RNA isolation and sequencing. **c)** *De novo* transcriptome assembly and annotation were conducted for each species. **d)** Differential gene expression analysis revealed the distinct immune response in the noble crayfish 3 days post-challenge with the Hap B strain of *A. astaci* and marbled crayfish 3 days post-challenge with the Hap A strain of *A. astaci.* Immune related DEGs were not present in either species 21 days post-challenge with *A. astaci.* **e)** Noble crayfish challenged with the Hap B strain of *A. astaci* were acutely infected and ultimately moribund, while the *A. astaci* Hap A challenged marbled crayfish showed high resistance to the pathogen, resulting infected without any mortality.

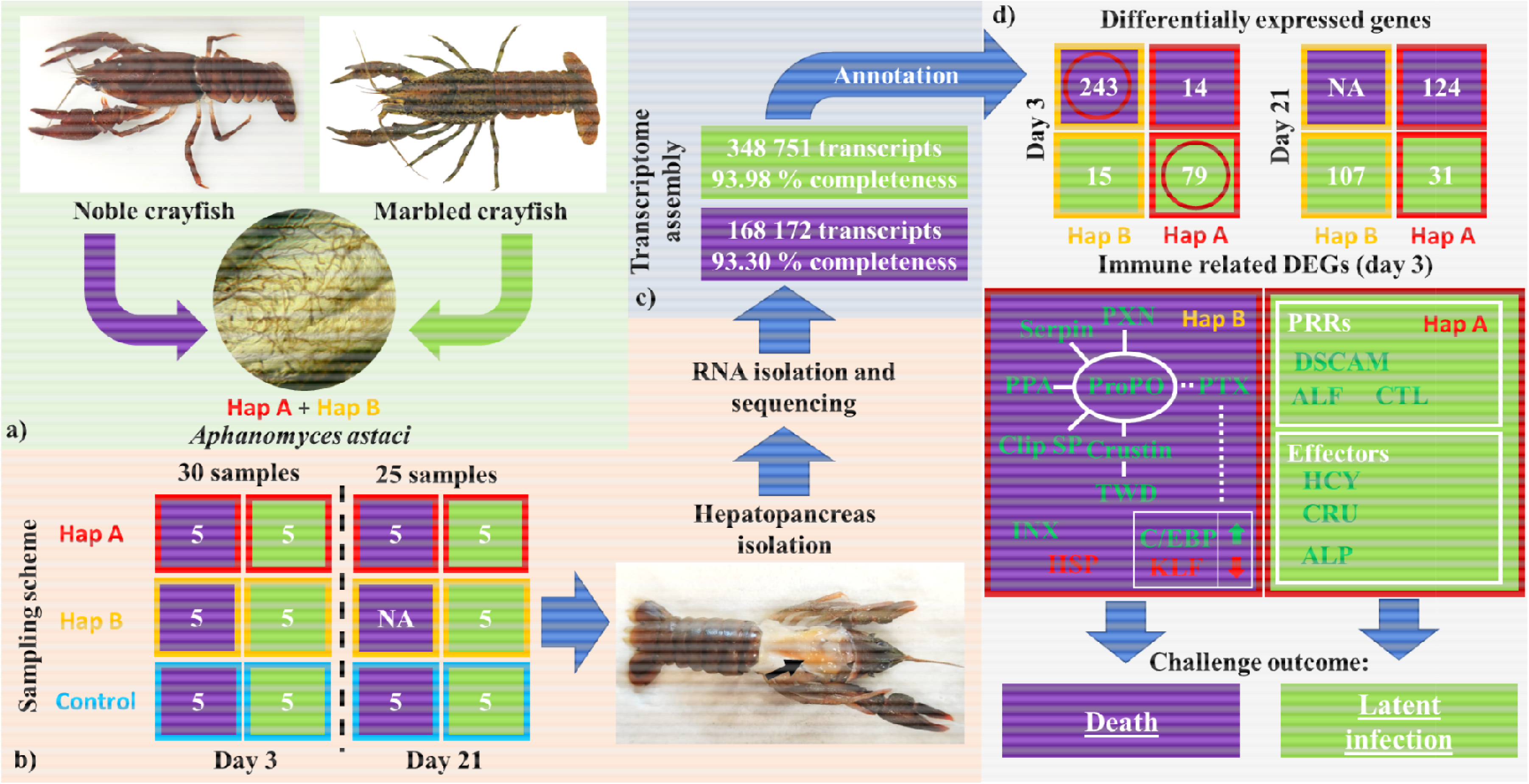

## Introduction

Freshwater crayfish are keystone species in freshwater ecosystems and are considered ecosystem engineers because of their ability to influence the trophic web and the freshwater habitat quality [1]. In the past two centuries, native European crayfish populations have faced a significant decline in number and size due to habitat loss, climate change and overfishing [2]. Introductions of highly competitive invasive crayfish species from North America represent one of the highest threats to the native freshwater crayfish species. North American crayfish act as carriers of the pathogen *Aphanomyces astaci*, the causative agent of crayfish plague disease [2]. This oomycete is causing mass mortalities and local extinctions among European crayfish populations [3]. Several North American crayfish species have so far established permanent populations across Europe, resulting in the presence of different *A. astaci* haplogroups of apparently variable within and between haplogroups virulence. The *A. astaci* strains present in Europe can be grouped into 4 different haplogroups [4]. To haplogroup A belong strains characterised by varying virulence, while haplogroups B, D and E are characterized by high virulence [5–7]. However, different crayfish species show diverse levels of susceptibility and resistance to varying strains of the pathogen.

North American crayfish species are generally considered resistant to the crayfish plague disease, likely because of their shared evolutionary history with *A. astaci* [8]. These crayfish are natural carriers of their specific *A. astaci* haplogroup, often efficiently preventing it from spreading inside their tissues through melanisation mediated encapsulation of the pathogen hyphae in the crayfish cuticle [9,10]. In contrast, European crayfish do not naturally carry the pathogen and are considered susceptible towards the disease, although resistant populations have been recently detected [7,11,12]. It has been hypothesised that one of the main factors contributing to the resistance of North American species is the constitutively over-expressed haemocyte prophenoloxidase (proPO), a key enzyme in the encapsulation of pathogens in melanin [13]. Conversely, in European crayfish the expression of this enzyme is dependent on stimuli of the pathogen [13]. The mechanisms underlying the crayfish immune response to *A. astaci*, however, is much more complex than the simple activation of the proPO cascade, but its molecular effectors and organs involved have not received much attention.

The immune response of crustaceans to pathogens comprises both cellular and humoral components, and the proPO cascade is only a small part of the humoral response [14–16]. Immune response in crustaceans is triggered by the pathogen associated molecular patterns (PAMPs), such as β-(1,3)-glucan, which is one of the main constituents of oomycetes cell wall [17]. These molecules are recognised by specific pattern recognition proteins (PRPs) of the host, which can exist as soluble molecules or as associated with cell membranes. PRPs of particular relevance are lectin-like proteins, Down Syndrome Cell Adhesion Molecules (DSCAMs) and Toll-like receptors (TLRs) [14,18]. The interaction between ligands and receptors leads to the activation of different molecular pathways involved in the humoral or cellular response, all of them coordinated by the core mediators of the crustacean immunity, the haemocytes. Haemocytes are crucial for the processes of phagocytosis, encapsulation and melanisation, and they are involved in delivering the molecular effectors of the humoral response, such as antimicrobial peptides and proPO, in the infection sites [16,19,20].

The hepatopancreas represents an integrated organ of the crustacean immunity and metabolism [21,22]. It plays a major role in pathogen clearance, antigen processing [23,24], detoxification, and heavy metal deposition [25]. It also serves as a source for immune molecules, which can be released from the epithelial cells into the haemocoel sinusoids, allowing for their rapid distribution in the haemolymph of the crayfish [22]. In recent years, the involvement of the hepatopancreas in the response to various disease and environmental factors has been highlighted in crustaceans [25–29]. However, its role in the immune response to *A. astaci* infection has not been clearly defined. Furthermore, the microbial community of the freshwater crayfish hepatopancreas remains unexplored.

In this study we aimed to deepen our understanding of the molecular mechanisms underlying the resistance and susceptibility of freshwater crayfish to the pathogen *A. astaci*. By analysing gene expression profiles of the hepatopancreas, we compared the immune response of the susceptible native European noble crayfish (*Astacus astacus*) and the resistant invasive marbled crayfish (*Procambarus virginalis*) to an *A. astaci* challenge. The marbled crayfish is a parthenogenetic freshwater crayfish species of North American origin, emerged after a triploidisation event in its closest relative *Procambarus fallax* from Florida [30,31]. Marbled crayfish is a known carrier of *A. astaci* [32] and is highly resistant to *A. astaci* infections [33]. In a controlled infection experiment, both species were infected with a highly virulent (haplogroup B, in further text Hap B) and a lowly virulent (haplogroup A, in further text Hap A) *A. astaci* strain [33]. The hepatopancreas of the crayfish was sampled three and 21 days post-challenge.

We hypothesised that the hepatopancreas is a highly relevant tissue in the immune response towards *A. astaci* infections, and we expected to detect several immunity related transcripts in all treatment groups. We expected that the gene expression profiles of the immune related transcripts differ among the noble crayfish and the marbled crayfish, reflecting the species' different ability to defend against the disease. Furthermore, for the susceptible noble crayfish we expected a stronger immune response when challenged with the highly virulent Hap B strain compared to the less virulent Hap A strain, while for the resistant marbled crayfish we did not expect any gene expression difference among treatment groups. Lastly, we expected the latently infected crayfish to show a chronic immune response against *A. astaci,* with the presence of differentially expressed immune related genes 21 days post-challenge.

The results presented in this paper deliver novel insights into the gene landscape involved in the immune response to the *A. astaci* challenge, deepening our understanding of the Crustacean immunity. The identification of genes responsible for higher resistance to *A. astaci* allows to screen species or populations to pinpoint vulnerable populations or populations with special interest for breeding purposes.

## 2. Materials and Methods

### 2.1. Experimental setup

A controlled infection experiment was previously conducted by Francesconi et al. [33] on marbled crayfish and noble crayfish. The crayfish were challenged with two different strains of *A. astaci*, a highly virulent Hap B strain and a lowly virulent Hap A strain. In total 55 individuals (30 marbled crayfish and 25 noble crayfish) were selected for RNA sequencing, with five replicates per treatment (Hap A, Hap B, control) from two time points (3d, 21d), with exception of the Hap B challenged noble crayfish group, where all crayfish were moribund in the first days of the challenge and were therefore all sampled in the first time point. For each individual a portion of the hepatopancreas was dissected and snap frozen in liquid nitrogen. For a detailed description of the experimental setup please refer to Francesconi et al., [33] and Boštjančić et al., [34].

### 2.2. Identification of the crayfish innate immunity genes and taxonomical distribution of transcripts

We retrieved a dataset of innate immunity related genes identified in Malacostraca by Lai and Aboobaker [35]. This dataset was expanded with the selected differentially expressed genes (DEGs) identified in the Hap B challenged noble crayfish. Furthermore, we included the genes specifically related to the proPO cascade. The complete list of used innate immunity genes and their respective sequences are available in the **Table S1 and File S1**. Transcriptome assemblies were queried against the subset of innate immunity related genes with BLASTn and BLASTx 2.10.1+. Hits were then inspected, their function was confirmed based on their e-value (lower than 1e-10), and the presence of the functionally important gene domains identified with a Pfam search.

The taxonomical distribution of the reads was reconstructed by conducting a DIAMOND 2.0.4 [36] search against the NCBI non-redundant protein database which contains sequences form GenPept, Swissprot, PIR, PDF, PDB, and NCBI RefSeq (sensitive mode, e-value≤1e-25). A single best BLAST hit was considered. The hierarchical distribution of BLASTx hit counts was further explored using interactive Krona 2.7.1 [37].

### 2.3. Read mapping

All sample reads were mapped to the newly obtained reference transcriptome [34] using the pseudo-alignment approach implemented in Salmon 0.13.1 [38]. Several “flags” were used in the Salmon mapping steps to correct the biases that might originate from sequence data: “-validateMappings” [39], “--seqBias” and “--gcBias” [40].

### 2.4. Differential gene expression analysis

Differential gene expression analysis was conducted according to the DESeq2 protocol [41] implemented in R with the following model design for noble crayfish: sex (male/female) + groups (Control vs Hap A or Hap B challenge) and for marbled crayfish: ~reproduction (yes/no) + groups (Control vs Hap A or Hap B challenge). Independent comparisons were conducted for each sampling point. Raw counts from the Salmon output were used as the input. Transcripts highly similar to the marbled crayfish and noble crayfish mitogenome, respectively, were removed prior to the analysis based on the BLAST hits against the mitogenome (NCBI accession number: KX279347.1 and NC_020021.1). Transcripts assigned to bacteria and archaea were also removed based on the DIAMOND search (see **2.2**). results Counts for individual Trinity transcript isoforms were grouped to Trinity genes with the tximport R package [42]. Lowly expressed genes were filtered out: only genes with the raw counts higher/equal to 10 across at least five samples were retained. The package “EnhancedVolcano” [43] was used for the visualisation of the DEGs and “apeglm” for noise removal [44]. The list of DEGs was exported and their counts, log2fold changes and adjusted p-values (FDR= 0.1, p-value= 0.05) together with their respective annotations were merged. Possible overlaps between the DEGs at different time points were inspected using Venn diagrams[45].

### 2.5. Gene set enrichment analysis

Enrichment of the innate immunity gene sets identified in the **2.2.** was conducted with ClusterProfiler [46]. Based on the results of the DESeq2 analysis, for each group all genes were ranked according to the following metric: −log10(x)/sign(y), where x is the p-value and y log2 fold change. To detect the enriched gene sets we used the GSEA() function, with the p values adjusted based on Benjamini-Hochberg correction for multiple testing (cutoff <0.01). Graphical representation of the results was obtained using the gseaplot2() function [46].

## 3. Results and discussion

With this study we compared the molecular immune response of the resistant marbled crayfish and the susceptible noble crayfish challenged with two *A. astaci* strains of different virulence, focusing on the gene expression profiles in the crayfish hepatopancreas. We investigated the relevance of the hepatopancreas in the anti-oomycete response, its capability of expressing a variety of immune related transcripts and contribution to the internal microbiome of the freshwater crayfish. We explore the landscape of differentially expressed genes in the response to *A. astaci* challenge and their relevance for the immune pathways, and we introduce several novel immune relevant molecules in freshwater crayfish. Furthermore, we integrate our results with the current knowledge on the host-pathogen coevolution between *A. astaci* and freshwater crayfish, indicating possible explanations for the observed differences between the species and across sampling groups. Lastly, we carefully consider the limitations of our approach, highlighting the necessary steps for future advancements in the field.

### 3.3. Hepatopancreas: mediator of the crayfish immune response to *A. astaci* challenge

#### 3.3.1. Innate immunity transcripts

In this study, we provide experimental evidence of the hepatopancreas involvement in mediation of the crayfish immune response to the *A. astaci* challenge through the synthesis of immune molecules. Activation of specific pathways and changes in the gene expression landscape are described in detail in **sections 3.4. and 3.5.** Here we provide an overview of genes involved in the immune related pathways and responses (**Figure 1**).

**Figure 1.**
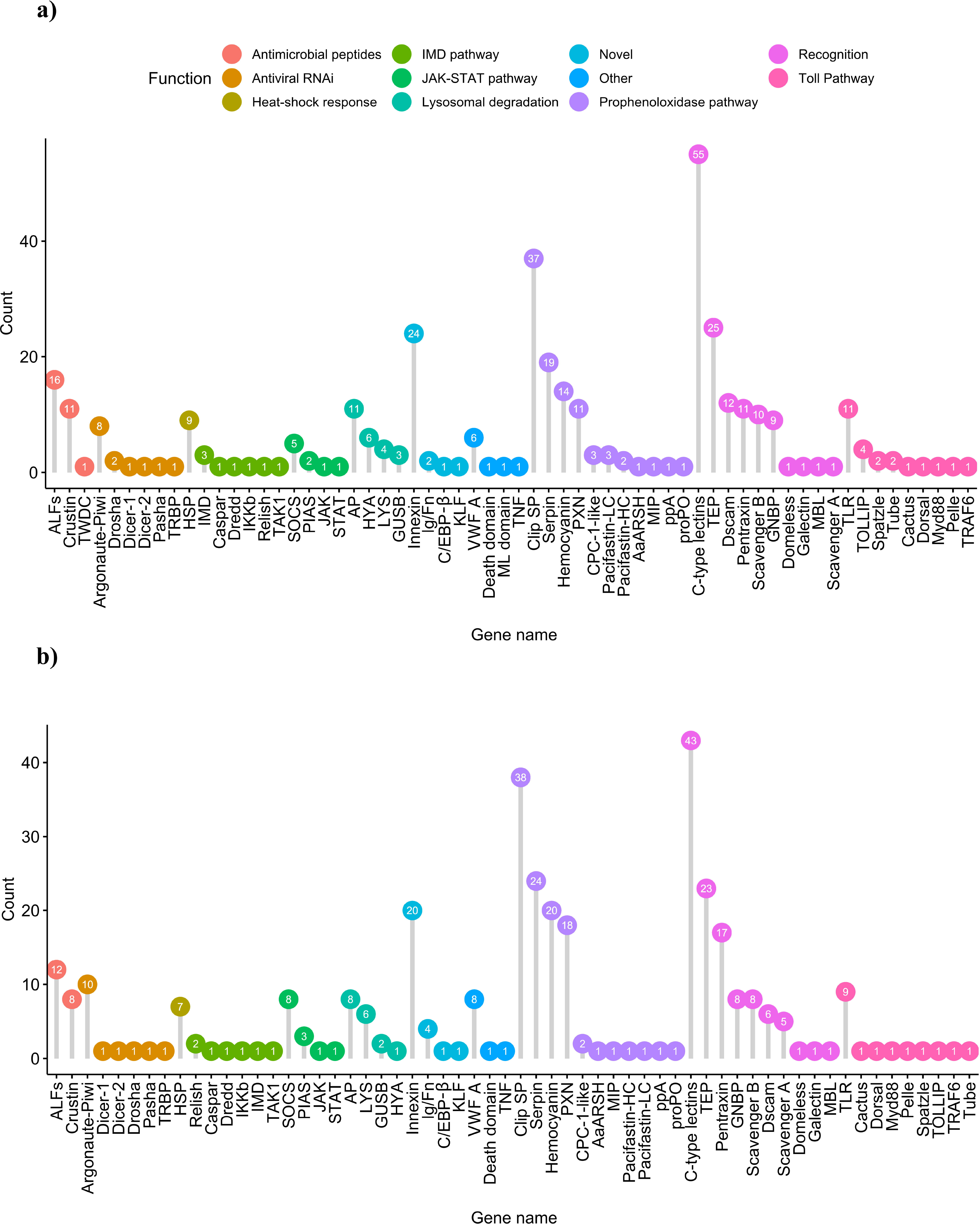
Genes involved in the representative immune related pathways, identified thought the similarity-based approach in **(a)** noble crayfish and **(b)** marbled crayfish. For all genes abbreviations are available in the Table S7.

Genomic research on non-model organisms is faced by the challenge of annotating large sets of genes from unknown origin. This challenge is particularly evident in Crustaceans [47,48], which are still largely underrepresented in genomic studies. To date, only 48 out of 727 genome assemblies representing Pancrustacea belong to Crustaceans (with the remaining 679 genomes belonging to Hexapoda) (Genomes-NCBI Datasets, accessed: April 2021). Furthermore, the canonical proPO pathway, considered a core immune response mechanism in the Crustaceans [49], is not represented in the KEGG database. Therefore, we conducted the annotation of the innate immunity related genes in the noble crayfish and marbled crayfish transcriptomes using a sequence and domain similarity-based approach. A total of 372 and 353 innate immunity related genes was identified through this approach in noble crayfish and marbled crayfish, respectively (**Figure 1**, **Table S2, Table S3, File S2, File S3**).

The identification of these innate immunity related genes provides a basis for future transcriptomic and genomic studies of the innate immunity in native and invasive freshwater crayfish species. For example, we successfully identified members of the immune signalling Toll pathway. This pathway is conserved in most members of Malacostraca [35]. In Hexapoda the activation of the Toll pathway is critical for antimicrobial peptides (AMPs) expression [50,51]. In the noble crayfish and marbled crayfish, we identified most of the Toll pathway-related genes as single copy (**Figure 1**). Recently, an extensive overview of innate immunity related genes has been conducted on numerous marine and freshwater Decapods [35]. The number of TLRs identified in those species ranged between 0 and 8, collocating the number of TLRs found in this study slightly above the higher value. Lastly, in the noble crayfish TOLLIP, Spätzle and Tube were detected in multiple copies (**Figure 1**).

The innate immune system in freshwater crayfish is armed with an arsenal of PRRs capable of recognising various PAMPs [52]. The β-(1,3)-glucan receptors (often referred to as Gram-negative binding proteins (GNBPs) or lipopolysaccharide binding proteins) play a vital role in the proPO cascade activation [53]. All GNBPs share a carbohydrate-binding β-glucanase domain as identified in this study [35]. The expansion of this family was previously reported in Decapoda [35], and confirmed in this study with 9 GNBPs identified in noble crayfish and 8 in marbled crayfish (**Figure 1**). Other molecules and pathways involved in the response to the *A. astaci* challenge are discussed in detail in the **section 3.5**.

#### 3.3.2. Hepatopancreas microbiome

In the taxonomical classification of the hepatopancreas transcriptome assemblies, 35,879 noble crayfish transcripts and 39,527 marbled crayfish transcripts had a significant BLASTx hit (**Figure 2)**. For both species, the majority of the transcripts had a significant hit among the eukaryotic taxa (66% for the noble crayfish and 68% for the marbled crayfish), among which the highest number of hits (51% and 52%, respectively) was assigned to the whiteleg shrimp (*Penaeus vannamei*). Large proportions of the BLASTx hits were assigned to bacterial taxa (26% and 23%, respectively). Among them, in the noble crayfish, the phylum Proteobacteria was represented with 24% of all hits, with the classes Gammaproteobacteria, contributing 9% of the total, Betaproteobacteria contributing 9% and Alphaproteobacteria contributing to 6% of all hits. In the marbled crayfish, the distribution of non-Eukaryotic hits was quite similar, with 18% Proteobacteria, and Gammaproteobacteria contributing 9%, Betaproteobacteria 6% and Alphaproteobacteria contributing 4% of all hits. From the Terrabacteria group, the class Actinobacteria represented 1% of all hits for noble crayfish, and 4% of all hits for marbled crayfish.

**Figure 2.**
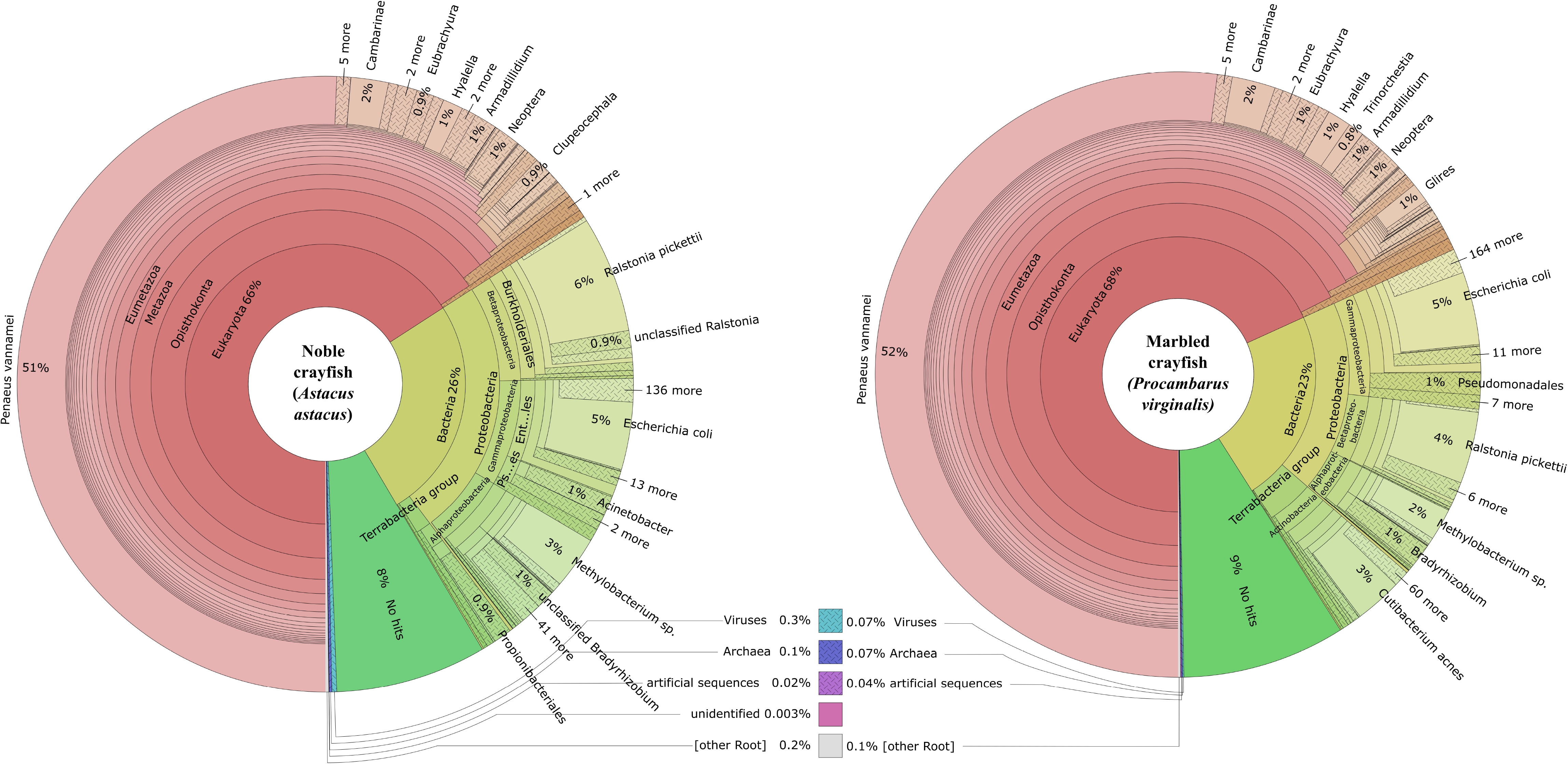
Krona plot summarizing the inferred taxonomies of assembled transcriptomes based on a Diamond search of the contigs against the non-redundant (nr) protein database.

The presence of bacterial communities in the hepatopancreas, as part of the crayfish digestive system, is not unexpected, and has been previously reported in other crustaceans like the giant tiger prawn (*Panaeus monodon*; [54]), the cherry shrimp (*Neocaridina denticulate*; [55]), and the whiteleg shrimp [56]. The phylum Proteobacteria seems to share a common predominance in the microbiome of Crustaceans [55]. Recently, a variety of bacterial species in the crayfish cuticle has been reported to contribute to the resistance of the signal crayfish and narrow-clawed crayfish (*Pontastacus leptodactylus*) to infections with *A. astaci* [57]. Our results revealed the signatures of some of these *A. astaci* growth-inhibitor bacteria within the hepatopancreas: *Pseudomonas chlororaphis* represented 0,8% and 0,5% of BLASTx hits in the noble crayfish and marbled crayfish, respectively, and *Acintobacter guillouiade* with 1% of BLASTx hits in noble crayfish. Further research, focused on the abundance of specific microbial taxa, is needed to explore a possible link between the *A. astaci* immune challenge and hepatopancreas microbial community composition. Nonetheless, it was proposed that microbiome communities may play a significant role in increasing host health, metabolism of protein and non-protein amino acids, as well as in modulation of the immune response and acting as competitors to the invading pathogens in crustaceans [58,59]. Furthermore, infection with pathogens (such as *A. astaci*) can cause microbiota dysbiosis [56]. Despite their obvious importance in metabolic and immunological functioning, microbiome communities of freshwater crayfish are mostly unreported in transcriptomic studies. Creating a knowledge database of the various microbial communities is necessary for understanding the immune response and disease progression in Crustaceans as well as in other Metazoans and might provide an effective tool in disease control [57].

### 3.4. Changes in gene expression profiles of *A. astaci* challenged crayfish

#### 3.4.1. Exploratory analysis of the mapping results

Mean mapping rate of the processed reads for the noble crayfish was 88.96% and for marbled crayfish 91.98% (**Table S4**). This was followed by the principal component analysis (PCA), performed to compare the replicates of the *A. astaci* exposed crayfish with the control group. The initial results of the PCA revealed a batch effect in noble crayfish and marbled crayfish samples (**Figure S1**). For the noble crayfish this effect was related to the differences between male and female individuals, accounting for 21% of the variance. For the marbled crayfish, the highest level of variance (63%) was caused by the differences between asexually reproducing and non-reproducing parthenogenetic females (see Francesconi et al. [33], for details). Therefore, in the down-stream differential gene expression analysis, we accounted for the sex of noble crayfish, as well as the reproductive status of marbled crayfish, by including them as factors in the DESeq2 analysis. After batch effect removal, the PCA analysis revealed the grouping only for the *A. astaci* Hap B challenged noble crayfish, while such grouping was revealed neither for other noble crayfish samples nor for the marbled crayfish (**Figure S1**).

#### 3.4.2. Differentially expressed genes

In the differential gene expression analysis, 35,300 genes for the noble crayfish and 52,491 genes for the marbled crayfish were analysed after the removal of the genes with low gene counts. In the noble crayfish, a total of 380 DEGs (202 up-regulated and 178 down-regulated) were detected in the response to challenge with *A. astaci* across all treatments (**Figure 3, Table S5**). The highest number of DEGs was observed in the Hap B challenged noble crayfish 3 days post-challenge, with 243 DEGs (141 up-regulated and 102 down-regulated) (**Figure 3**), with many involved in the immune response (**Figure 4**). The lowest amount of DEGs was observed in the Hap A challenged noble crayfish 3 days post-challenge, with only 14 DEGs (7 up-regulated and 7 down-regulated) (**Figure 3**). The DEGs relevant to the innate immunity, mainly connected to the proPO cascade, were observed in the Hap B challenged noble crayfish 3 days post-challenge (**Figure 3).** In the marbled crayfish a total of 232 DEGs (102 up-regulated and 130 down-regulated) were detected in the response to the challenge with *A. astaci* across all treatments (**Figure 3, Table S6**). The highest number of DEGs related the innate immunity was observed in the Hap A challenged marbled crayfish 3 days post-challenge, with 79 DEGs (47 up-regulated and 32 down-regulated), and the highest overall number of DEGs in the marbled crayfish was observed 21 days post challenge with the Hap B strain 107 DEGs (40 up-regulated and 67 down-regulated). The lowest amount of DEGs was observed in the Hap B challenged marbled crayfish 3 days post-challenge, with only 15 DEGs, all down-regulated (**Figure 3, Table S6**).

**Figure 3.**
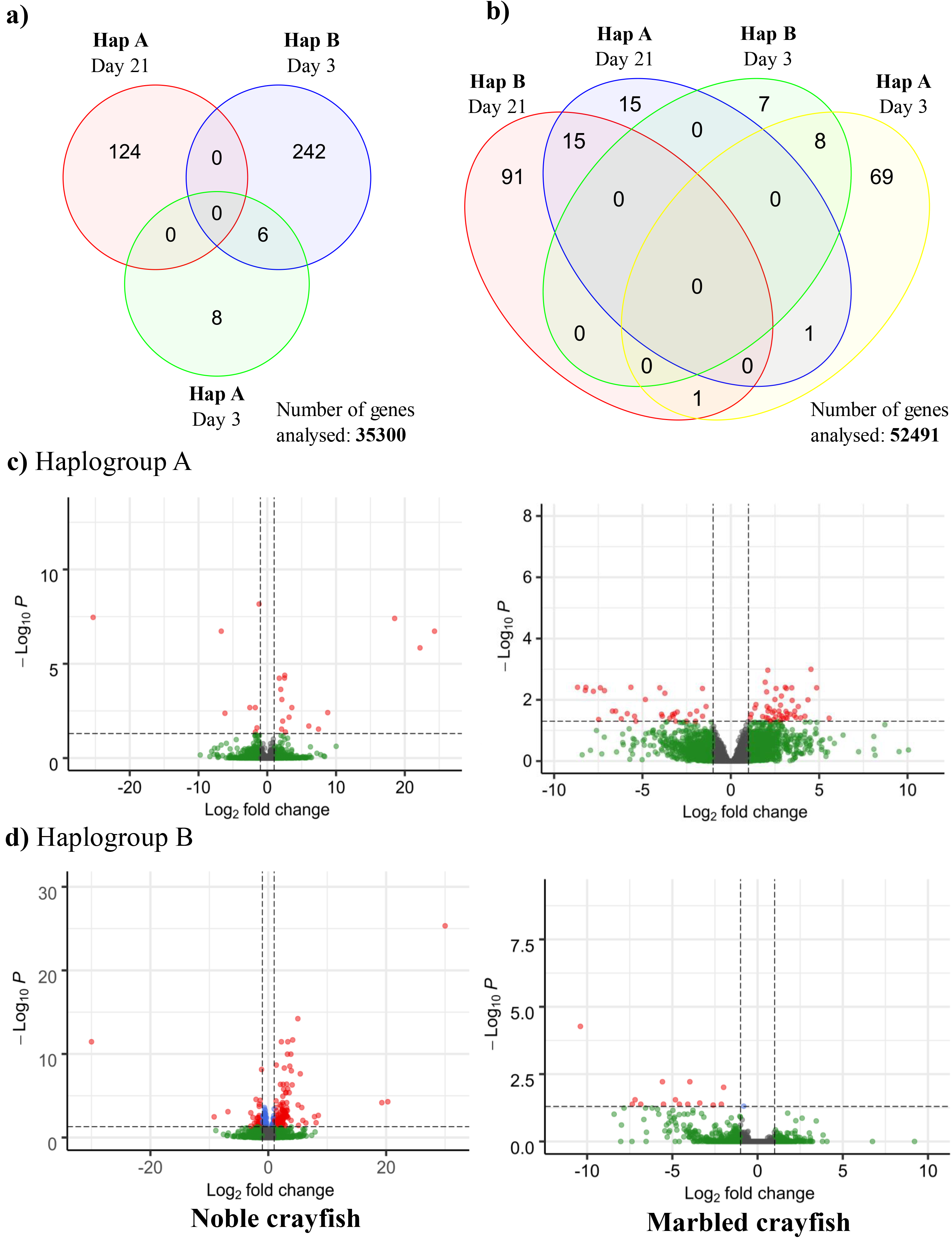
Results of the differential gene expression analysis. **(a)** Venn diagram representing DEGs for all treatments in the noble crayfish **(b)** Venn diagram representing differentially expressed DEGs for all treatments in the marbled crayfish. Volcano plots for the noble and marbled crayfish. **(c)** 3 days post-challenge with haplotype A, **(d)** 3 days post-challenge with haplotype B. The threshold values are represented as dashed lines (p-value = 0.05, Fold change = 2). Genes above fold change and p-value threshold are coloured red.

**Figure 4.**
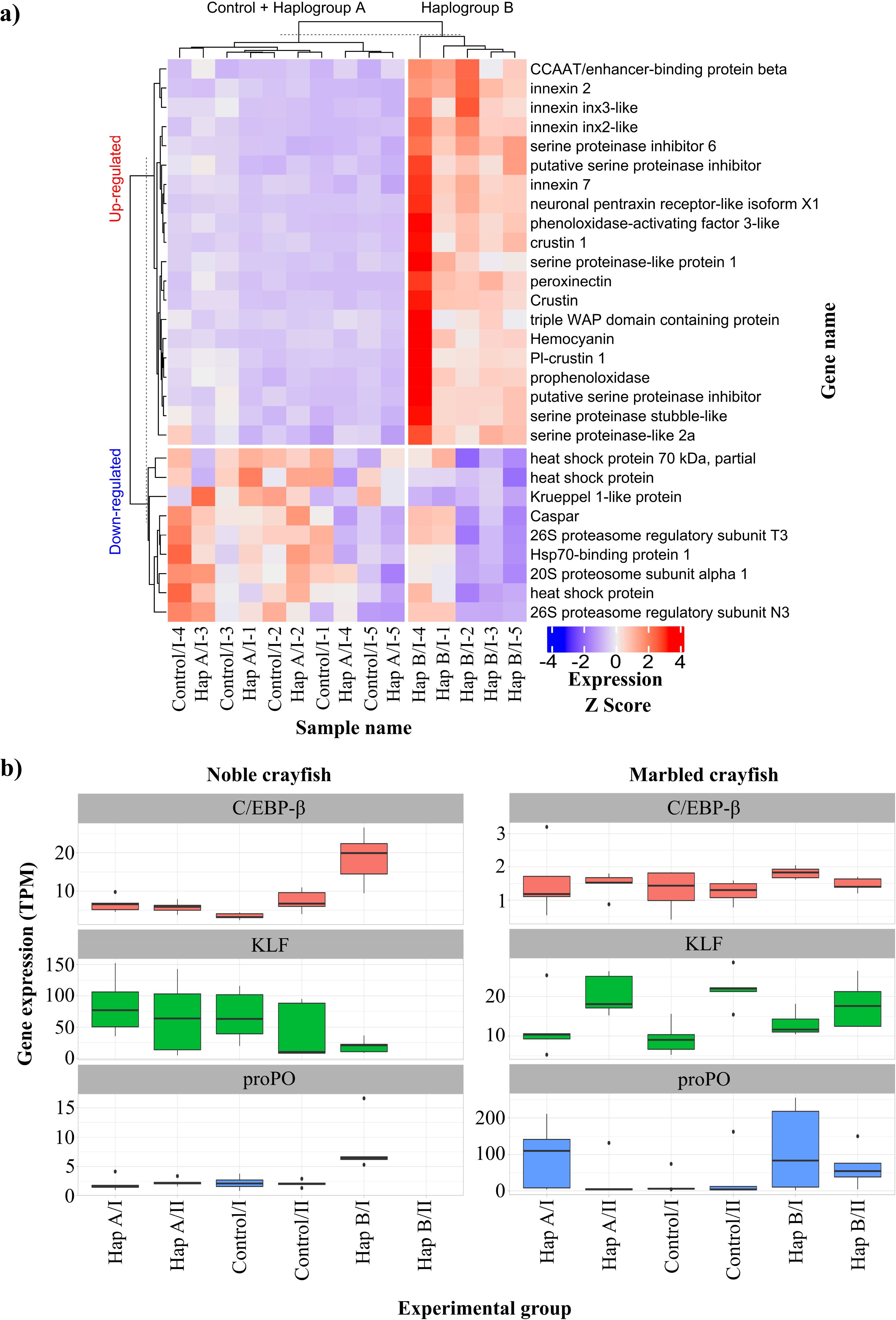
Heatmap of the immunity genes for each sample and treatment detected as differentially expressed in the noble crayfish **(a)** Raw counts were transformed to transcripts per million (TPM), followed by standardisation with Z-score scaling (where Z score is calculated as follows: Z= s_i_-μ/σ where s_i_ is the gene expression for a sample in TPM, μ is mean of the expression for each gene in TPM and σ is standard deviation of the expression for each gene in TPM). Therefore, the colours in the heatmap reflect the relative expression levels between samples per each gene, with higher expression in red and lower expression in blue. Hap A, haplogroup A; Hap B, haplogroup B, I and II, first and second sampling point, respectively (3 days and 21 days post-challenge), 1-5, identifying number of the crayfish **(b)** gene expression of the prophenoloxidase (proPO), CCAAT/enhancer-binding protein beta (EBP), and Krueppel like protein (KLP) in the marbled crayfish and noble crayfish challenged with *A. astaci*. Expression values are shown in TPM.

Our results indicate the absence of a chronic immune response to the challenge with *A. astaci* in both species. The lack of the highly differentially expressed immune related genes 21 days post-challenge with *A. astaci* suggests that the active immune response in the hepatopancreas had already come to an end, or was capped below the detection level of the differential gene expression analysis at the time of the second sampling (see 3.4.3). However, a chronic response could be mediated, as previously suggested, by circulating haemocytes in the haemolymph of latently infected crayfish [60]. Future studies focused on different immune system relevant tissues in crayfish, such as gills or haemocytes, might clarify this aspect.

#### 3.4.3. Enriched gene sets in the response to the A. astaci challenge

As a complementary approach to the differential gene expression analysis, we utilised the newly identified immunity genes (see section 3.3.1.) to conduct a gene set enrichment analysis. This approach allowed us to detect moderate or minor changes in the gene expression data [61]. For the noble crayfish, our results revealed the enrichment of AMP, proPO pathway and novel (encompassing novel genes identified in this study) gene sets in the Hap B challenge group (**Figure 5**) and recognition gene set in the Hap A group 21 days post-challenge (**Figure S2**). The proPO pathway gene set was under-represented in the Hap A challenged noble crayfish 3 days post-challenge. In the marbled crayfish, AMP, proPO and recognition gene sets were enriched for the Hap B challenged group at both sampling points (**Figure S2**). Furthermore, in the Hap A challenged group, recognition and proPO gene sets were enriched (**Figure 5**). In the marbled crayfish, 21 days post-challenge with Hap A we detected no enriched gene sets. These results, in line with the differential gene expression analysis, suggest that proPO pathway, AMPs and recognition proteins, although not detected as differentially expressed, play a major role in the response to the *A. astaci* challenge. Their interplay and significance are discussed in the further text.

**Figure 5.**
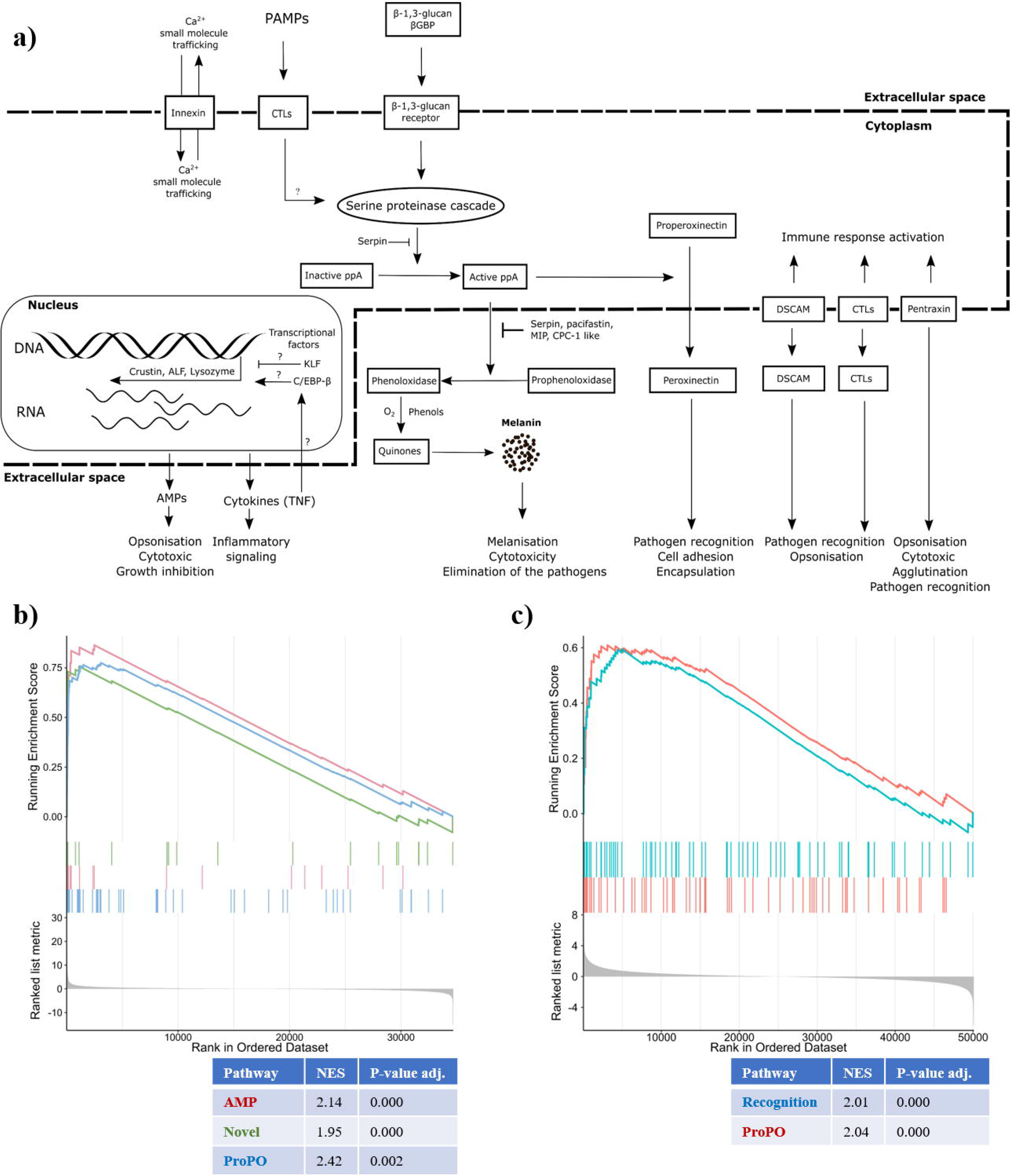
Pathways involved in the freshwater crayfish immune response to *A. astaci* immune challenge, (a) Schematic representation of the crayfish immune response to *A. astaci* challenge (b) Results of the gene set enrichment analysis for the noble crayfish challenged with Hap B strain of *A. astaci* (Day 3), (c) results of the gene set enrichment analysis for the marbled crayfish challenged with Hap A strain of *A. astaci* (Day 3)

### 3.5. Molecular mechanisms of the immune response to the *A. astaci* challenge

#### 3.5.1. Activation of prophenoloxidase cascade

In our study, we observed an up-regulation of proPO, serine protease (ppA) and peroxinectin (PXN) in the hepatopancreas of the Hap B challenged noble crayfish (**Figure 4**). The activation of proPO cascade is the most recognised humoral response among crustaceans (**Figure 5**) [53,62]. Phenoloxidase (PO) synthesized in its zymogen/inactive form (proPO), is the central enzyme of the pathway. It is cleaved by its activating ppA to the catalytically active PO and the 20 kDA N-terminal fragment, ppA-proPO, with a strong agglutination and bacterial killing capacity [63]. Activated PO is involved in the conversion of the phenolic substances into the toxic quinone intermediates involved in the production of melanin, the terminal pathogen encapsulating agent of the proPO cascade [60]. Alongside PO, ppA activates the formation of PXN, involved in pathogen recognition, cell adhesion and encapsulation [64,65]. It was previously assumed, that only the mature haemocytes (granular and semigranular), which are responsible for the release of the proPO in the response to the pathogen stimulation [49,62], are characterised by the onset of proPO expression [15]. Our results suggest that hepatopancreas is involved in the production of the central proteins of this pathway (**Figure 4**).

We observed that the challenge with *A. astaci* caused an up-regulation of proPO only in the Hap B challenged noble crayfish (**Figure 4**). Previously, differences between the expression levels of proPO have also been observed in *A. astaci* susceptible and resistant crayfish [13]. Specifically, it was proposed that the proPO expression is continuously elevated in the invasive signal crayfish and is non-responsive to immune stimuli with the β-1,3-glucan polysaccharide (laminarin). On the other hand, in the susceptible noble crayfish, proPO expression is constitutively at lower levels, although it can be elevated with the injection of laminarin. It should be noted that one of the main constituents of the oomycete cell wall is β-(1,3)-glucan, which binds to the specific GNBP located on the haemocyte cell membrane [17]. The GNBPs play an essential role in the activation of proPO cascade [66]. Our findings indicate the expression of proPO in both susceptible and resistant crayfish can be altered in response to pathogen stimulation. Moreover, the variances in the proPO expression levels (transcripts per million, TPM) were much higher in the marbled crayfish challenged with Hap A of *A. astaci* 3 days post-challenge and Hap B of *A. astaci* three- and 21-days post-challenge, than in the noble crayfish challenged with Hap B of *A. astaci.* (**Figure 4**), which was also complemented with the results of the GSEA (**Figure 5, Figure S2**)

Our results indicate that in the Hap B challenged noble crayfish, several serine proteinases (Clip SPs) and serine proteinase inhibitors (serpins) were up-regulated in the response to the infection (**Figure 4, Table S5**), and pacifastin-HC protein was up-regulated in the Hap A challenged marbled crayfish 3 days post-challenge (**Table S6**). These proteins are responsible for the spatial and temporal control of the proPO cascade (**Figure 5**) [60]. Excessive activation of the proPO pathway can cause damage to the host due to the production and release of toxic quinones, therefore inhibitory proteins are of utmost importance. In particular, the proteins involved in the proPO regulation are: pacifastin, a regulatory inhibitor of ppA [67]; melanisation inhibition protein (MIP) [68]; caspase 1-like molecule (CPC-1-like), released concomitantly with the proPO limits the proteolysis of proPO; and mannose-binding lectins [63]. Serpins were reported to play a role in the proPO cascade inhibition [69]. The recognition of the oomycete β-(1,3)-glucan activates the Clip SP cascade responsible for cleavage of the ppA [49]. The up-regulated serpins could also be involved in the inhibition of the oomycete proteinases [70]. Thus, serpins exhibit a dual role as an anti-oomycete agent, as well as the protectors against the proPO cascade overactivation [22,71]. This is further supported by the high number of genes encoding for the putative Clip SP (37 in the noble crayfish and 38 in the marbled crayfish) and their inhibitor serpins (19 in the noble crayfish and 24 in the marbled crayfish). The expansion of the Clip SP in Malacostraca (compared to the other Pancrustacea) was previously observed by Lai and Aboobaker [35] with the highest number of Clip SP (72) observed in the whiteleg shrimp. Co-expression of the proPO cascade effectors, and the proPO inhibitors in the hepatopancreas of Hap B infected noble crayfish, suggests that the proPO cascade is highly involved in the response to the *A. astaci* challenge.

Although only one gene was annotated as the putative proPO, multiple hemocyanin (HCY) domain containing genes (14 in noble crayfish and 20 in marbled crayfish) were uncovered in both species (**Figure 1**). HCY is evolutionarily closely related, but distinct, to proPO [72]. It is believed that Crustacean HCYs can, to a certain extent, mimic the proPO functions [49]. Crustacean HCY is a large type-3 copper containing respiratory protein which forms hexameric structures responsible for oxygen transport [73]. Alongside proPO, in the Hap B challenged noble crayfish, one of the HCY containing proteins was observed as up-regulated (**Figure 4, Table S5**). In the marbled crayfish challenged with the Hap A 3 days post-challenge, a highly expressed HCY containing protein was also observed as up-regulated in the hepatopancreas (**Table S6**). Unlike vertebrate hemoglobins, HCYs are cell-independent, and are solely suspended in the crayfish haemolymph [73]. This means that the HCYs can be directly excreted from the hepatopancreas, where they are synthesised, to the crayfish haemolymph, without damage to the organism [74,75]. On the other hand, proPO must be transported to the infection site and incorporated in the granules of semi-granular and granular haemocytes (blood cells) [13,60]. Shortly after the immune challenge, a significant drop in the number of circulating haemocytes (condition termed haemocytopenia) is observed due to haemocyte mobilisation to the infection site [20,76]. These haemocytes are mainly directly replaced during haematopoiesis from the hematopoietic tissues [17]. This usually occurs 12-48 hours after the initial challenge to the innate immunity [16,76]. Therefore, during the period of circulating haemocyte depletion, both sensitive and resistant crayfish can rely on the components of the humoral innate immune response, such as antimicrobial peptides and HCYs, until the haemocyte replenishment. This is concordant with the observation by Decker et al., [73] that suggests the innate immunity involvement of the high concentration of HCYs in the circulating haemolymph in tarantula [77]. Lastly, HCYs can be proteolytically processed, resulting in a release of AMPs, such as those belonging to the astacidin family [78].

#### 3.5.2. Expression of pattern recognition receptors (PRRs)

We observed two up-regulated putative C-type lectins (CTLs) in the marbled crayfish, one in the *A. astaci* Hap A challenged group 3 days post-challenge and one in the *A. astaci* Hap B challenged group 21 days post-challenge (**Table S5**). Lectins are a diverse group of proteins capable of binding carbohydrate-binding domains with high specificity [79]. In crustaceans, lectin recognition leads to downstream activation of cellular and humoral responses such as agglutination [80], endocytosis [81], encapsulation and nodule formation [82], synthesis of AMPs [83], antiviral activities [84], and melanisation through the proPO cascade activation [85]. We have identified 55 putative CTLs in noble crayfish and 43 putative CTLs in marbled crayfish (**Figure 1**). Among PRRs, CTLs have a major role in the innate immunity of freshwater crayfish, where they have also experienced a major increase in their diversity [35].

Among the differentially expressed genes involved in pattern recognition we observed an up-regulated DSCAM in the marbled crayfish challenged with *A. astaci* Hap A 3 days post-challenge (**Table S6**). DSCAM is a member of the immunoglobulin (Ig) superfamily, with a similar structure in both mammalians and invertebrates. The DSCAM molecule consists of three main components, an extracellular region with several Ig and fibronectin type III domains, a transmembrane domain, and a cytoplasmic tail. Unlike its mammalian counterpart, invertebrate DSCAM exhibits hypervariability in the extracellular domains achieved through a mechanism of alternative splicing during mRNA maturation [86,87]. In total, we identified 12 putative DSCAM-encoding genes in the noble crayfish and 6 in the marbled crayfish (**Figure 1**). DSCAM molecules have been shown to be involved in the antiviral [88] and antibacterial response, mainly in the opsonisation [53]. It is worth noting that due to their hypervariable domain, Dscams are considered likely key molecules for immunological memory in crustaceans [18]. Both CTLs and DSCAMS can exist in a membrane bound and secreted form [89,90]. Therefore, CTLs and DSCAMS expressed in the hepatopancreas of crayfish can probably be excreted directly to the haemolymph upon the immune challenge, acting as a part of the humoral immune response mechanisms to the pathogen infection.

Alongside DSCAM we observed another immunoglobulin/fibronectin (Ig/Fn) domain containing protein up-regulated in the marbled crayfish challenged with *A. astaci* Hap A 3 days post-challenge (**Table S6**). This protein shared 27% identity with the fruit fly (*Drosophila melanogast*er) protein amalgam (Ama, NCBI acc. No.: P15364.2). This amalgam-like protein was 510 amino acid (aa) long, with a molecular weight of 55.63 kDa. It contained 1-21 aa signal peptide domain, three Ig domains (67-158 aa, 166-254 aa, 257-345 aa), and a Fn domain (347-453 aa) with a cytokine receptor motive (439-443 aa). In total, we identified 2 Ig/Fn domain containing proteins with this domain organisation in the noble crayfish and 4 in the marbled crayfish (**Figure 1**). The presence of the C-terminal Fn domain clearly distinguishes this protein form the fruit fly Ama [91]. Nonetheless, we can hypothesise that this protein could share the secreted nature of Ama, and its cell adhesion properties [92], potentially having a role in opsonisation, and immune response mediation through its cytokine receptor motive located in the fibronectin domain.

Among the up-regulated DEGs in the Hap B challenged noble crayfish, we identified a pentraxin domain containing gene (**Table S5**, Pfam: PF00354). The protein product of this gene is 254 aa long (27.95 kDa), with a signal peptide (1-21 aa) on the N-terminus and only 55.51% identity with the neuronal pentraxin receptor-like isoform X2 from the whiteleg shrimp (XP_027224174.1, identified with Blastx). Like the most-well studied pentraxins, (e.g. C-reactive protein (CRP) or Serum-amyloid P component (SAP)), due to its size this pentraxin probably belongs to the group of short pentraxins [93]. We identified 11 putative pentraxin genes in the noble crayfish and 17 in the marbled crayfish (**Figure 1**). Pentraxins (or pentaxins) represent a multifunctional and evolutionary conserved group of proteins, with a critical role in the humoral innate immune response [94]. They can recognise a wide range of the pathogen associated molecular patterns, and serve as opsonin, cytotoxic effectors, agglutination promotors or as activators of the complement [93,95,96]. Not much is known about the complex system of the complement in the freshwater crayfish and previously hypothesised pentraxin complement activation is most likely not mediated through the C3 component of the complement [95], as it is in vertebrates [96] since C3-like proteins have reportedly been lost in Pancrustacea [35].

In endothermic animals the source of pentraxins is the liver [97] and in the horseshoe crab (*Limulus polyphemus*) and American lobster (*Homarus americanus*) these proteins are produced in hepatopancreas [98,99]. From there they are released to the haemolymph. Pentraxins are classical acute phase proteins. In humans, CRP can be utilised as a marker of bacterial and fungal diseases progression [95]. To our knowledge, this is the first time a pentraxin-domain containing protein is identified in crayfish in the response to *A. astaci* infection. This acute protein could be a good indicator of the disease progression. Application of the acute phase proteins as the markers of the immune status has been previously proposed for the American lobster, where pentraxin-domain containing protein has been recognised as an important component of the immune response to the pathogen challenge [47,99,100]. Involvement of the recognition proteins in the response to the *A. astaci* challenge was further supported by the results of the GSEA (**Figure 5, Figure S2**).

#### 3.5.3. Antimicrobial peptides as effectors of the innate immune response

In the noble crayfish challenged with the Hap B strain we identified three up-regulated crustins (**Table S5**). Among them, of particular interest was the DE triple whey acidic protein (TWP) domain containing crustin, identified in the noble crayfish but with no ortholog in the marbled crayfish. In the noble crayfish we identified 11 and in marbled crayfish eight putative crustins (**Figure 1**). Crustins are part of the cationic antimicrobial peptides AMPs and have three main components: the signal peptide, the multi domain region at the N-terminus and the whey acidic protein (WAP) domain at the C-terminus. They are classified in five groups based on their structure (type I-V) [101]. Crustins are mainly expressed in the crayfish haemocytes, where they can be rapidly secreted directly into the haemolymph during the immune challenge [102,103]. Some crustins can also exhibit antiprotease activity, possibly inhibiting the proteases secreted by *A. astaci*, limiting the pathogen growth [104]. Recently, a novel TWD containing crustin was described in the red swamp crayfish (*Procambarus clarkii*), showing antibacterial activity [105]. In the marbled crayfish challenged with the Hap B strain we identified one up-regulated crustin 21 days post-challenge (**Table S6**). Crustins may play an important role in the anti-oomycete response of the freshwater crayfish and require a closer attention in future. TWD containing crustins might be of special interest, due to their presumed tissue wide expression profiles and participation in the host immunity throughout the whole body [105].

Up-regulated antilipopolysaccharide factor (ALF) was identified in the Hap A challenged marbled crayfish 3 days post-challenge (**Table S6**), while DE ALFs were not detected in the noble crayfish. This suggests that ALF up-regulation might play a vital role in the resistance of the marbled crayfish towards the *A. astaci* challenge, possibly by binding to the oomycete β-1-3-glucan, hence increasing the host antimicrobial defences acting as an opsonin for the haemocytes [101]. In the noble crayfish, we identified 16 putative ALFs, and in the marbled crayfish we identified 12 putative ALFs (**Figure 1**). ALFs are small proteins with the hydrophobic N-terminal region forming, three β-sheets and three α-helices [35], Pfam: DUF3254. They have been observed in the wide range of crustaceans [106], and they are expressed in a wide range of tissues, showing growth inhibiting activity towards bacterial and fungal microorganisms, as well as opsonic activities [107,108]. Like crustins, they possess a signal peptide domain and can be excreted [101]. AMPs were enriched in both noble crayfish and marbled crayfish challenged with Hap B strain (**Figure 5, Figure S2**).

#### 3.5.4. Innexins: involvement of the gap junction proteins in the crayfish innate immunity

Among the differentially expressed genes, we detected four up-regulated innexins (INXs) in the Hap B challenged noble crayfish 3 days post-challenge (**Table S5**). These proteins represent the subunits that compose the hemichannel of the gap junctions, and they are analogous to the vertebrate connexin subunits [109]. Gap junctions represent the sites of the direct cell to cell communications. This interaction is achieved through the formation of the plasma membrane spanning channels, with each cell contributing to one half of the channel. The mechanisms of gap-junction communications and their repercussions have long been studied in vertebrates, where they are widely distributed across tissues [110,111]. Although these channels were first observed in the 1950s in the noble crayfish cells, their involvement in the immunity of freshwater crayfish species is not well understood [112]. We identified 23 putative INXs in the noble crayfish and 20 putative INXs in the marbled crayfish (**Figure 1**). For comparison, 8 INXs were identified in the fruit fly, 25 INXs in the roundworm (*Cenorabditis elegans*), 21 in the mediterranean medicinal leech (*Hirudo verbana*) and 6 in the Jonah crab (*Cancer borealis*) [113–116]. In the mud crab (*Scylla paramamosin*), Sp-inx2 expression was up-regulated in the hepatopancreas, gills and haemocytes after challenge with bacteria, and was highly expressed in the haemocytes under normal conditions [117]. Although the roles of INXs in invertebrates are largely unknown, based on the current knowledge of the functions of gap junction proteins in other species, we can argue that they could be involved in the antigen processing, as well as in the metabolic and signalling molecules trafficking [118]. This further establishes the role of the hepatopancreas as a key organ in the distribution of the immune molecules to the crayfish haemolymph [22]. Further studies are needed to elucidate the roles of INXs in invertebrate immunity.

#### 3.5.5. Transcriptional factors as novel components in the response to A. astaci challenge

Changes in the gene expression levels are controlled through a set of specific transcription factors that interact with the gene regulatory sequences, present in the promoter and enhancer regions. In the Hap B challenged noble crayfish 3 days post-challenge we identified both up-regulated and down-regulated genes, which serve as transcription factors and *bona fide* play vital roles in the immune response to the pathogen (**Table S5**). One of these genes is a master gene expression regulator belonging to the CCAAT/enhancer-binding protein (C/EBP) family [119]. This family is involved in the regulation of cellular growth, differentiation and death, as well as in haematopoiesis, and immune and inflammatory processes during various diseases [119,120]. The expression of the putative CCAAT/enhancer-binding protein beta (C/EBP-β), present in single copy in both noble crayfish and marbled crayfish, was up-regulated in the noble crayfish challenged with Hap B, while the expression levels in marbled crayfish remained unchanged (**Figure 1, Figure 4**). It has been shown that the expression of the ALFm3 (member of antilipopolysaccharide factor family) in the giant tiger prawn is under the control of C/EBP-β [121]. Previously it has also been shown that C/EBP-β binding sites are present in the crustin Pm7 [122]. The interaction of the C/EBP-β and NF-κB, key transcriptional factor in Toll and IMD pathways was reported during the promotion of the inflammatory mediator’s gene expression [123]. In mice, C/EBP-β is responsible for the control of tumor necrosis factor alpha (TNFα), SAP, complement C3 component expression [119]. This could suggest that the putative C/EBP-β up-regulation is crucial for the acute phase of the *A. astaci* infection in the noble crayfish.

Furthermore, we detected a down-regulation of putative Krüppel 1-like factor protein (KLF1), a member of the Krüppel-like factor (KLF) family, in the noble crayfish challenged with *A. astaci* Hap B (**Table S5, Figure 4**). Members of KLF family are transcription factors involved in a variety of metabolic pathways and in the energetic homeostasis of various tissues [124]. KLF1 belongs to a group of KLFs which function primarily as transcriptional activators, although interaction with transcriptional repressors has also been reported [124]. It is present in single copy in both noble crayfish and marbled crayfish (**Figure 1**). In humans, KLF4 is heavily implicated in the regulation of the anti-fungal response to *Aspergillus fumigatus* and *Candida albicans* and was identified as the only transcriptional factor down-regulated during the immune challenge [125]. It has been shown that in whiteleg shrimp, the host LvKLF is important for the replication and gene expression of the viral pathogen [126,127]. In the giant river prawn (*Macrobrachium rosenbergii*), it has been shown that MrKLF is an important regulator of expression of four antimicrobial peptides, namely Crustin (Crus) 2, Crus8, ALF1, and ALF3 [128]. Knowledge on the expression and regulation of invertebrates KLF is lacking, therefore conclusive interpretations for the function of the putative KLF1 require further research efforts. Based on the change in the KLF1 expression levels in noble crayfish, we might speculate that KLF1 repression is important for the activation of the immune response genes in this species. In the marbled crayfish KLF1 expression levels are unchanged during *A. astaci* challenge (**Figure 4**).

Together with KLF1 we also detected down-regulation of Caspar, a transcriptional suppressor homologous to the Fas-associating factor 1, in the noble crayfish challenged with *A. astaci* Hap B (**Table S5, Figure 4**). This transcriptional factor has been shown to play a critical role in the fruit fly, negatively affecting its antibacterial resistance through inhibition of the IMD pathway [129]. In both species Caspar was detected in a single copy (**Figure 1**).

#### 3.5.6. Other DEGs in the response to A. astaci challenge

Among the up-regulated DEGs in the marbled crayfish we observed several other immune related genes, such as Tumour necrosis factor (TNF) domain-containing protein (Panther entry: PTHR15151; protein Eiger; putative cytokine) and lysosomal enzyme putative alkaline phosphatase (AP) (**Table S6**). Cytokines, such as TNFs are heavily involved in the mediation of the immune and inflammatory responses [130]. They are also known activators of the extracellular trap release (ETosis), a microbicidal mechanism [131]. TNF is also a downstream target of the above mentioned KLFs [125]. Moreover, in the fruit fly, TNF homolog Eiger is responsible for the release of proPO in the crystal cells [132]. TNF is also an activator of the C/EBPβ expression and DNA binding activity [120]. The implication of this gene in the regulation of anti-oomycete responses remains to be experimentally proven in future studies. Alkaline phosphatase, β-glucuronidase, lysozyme, esterases and proteases have been recognised as some of the main lysosomal enzymes in the invertebrates [20]. Lysosomal activity has been implicated in the mechanism of antigen processing in the hepatopancreas epithelial cells and their subsequent release into the haemolymph in giant tiger prawn [22,23,133]. This observation might further establish the role of hepatopancreas in building the immune tolerance to the *A. astaci* challenge.

Interestingly, we uncovered four members of the heat-shock protein (HSP) family (HSP70-like, HSP-like-1, HSP-like2 and HSPBP 1) together with proteasome components (20S proteosome subunit alpha 1, 26S proteasome regulatory subunit N3 and 26S proteasome regulatory subunit T3), as down-regulated in the acutely infected noble crayfish, 3 days post challenge with Hap B strain (**Table S5, Figure 4**). Establishing a correct protein conformation is important for the protein activity. Failure to do so could be due to a lack of molecular chaperons, such as members of the HSP family [134]. Moreover, down-regulation of the ubiquitin mediated proteolysis proteasome genes might have led to the misfolded protein aggregation. It has been shown that HSP 70 is up-regulated in the anti-viral response to the White spot syndrome virus (WSSV) in the giant tiger prawn [135] and the red swamp crayfish [136]. In the fruit fly, it has been shown that HSP 27 has an antiapoptotic activity, inhibiting the TNF-mediated cell death [137]. This might suggest that during the *A. astaci* challenge, in acutely infected noble crayfish, a tissue wide apoptosis is in progress.

### 3.6. Coevolutionary aspects of the host immune response to the pathogen challenge

Our experimental setup allowed us to characterise and compare the immune response of the noble crayfish and marbled crayfish challenged with *A. astaci*. By challenging both crayfish species with two different *A. astaci* strains of different origin and virulence, we can make inferences on coevolutionary aspects of the host immune response to the pathogen challenge. The utilized Hap B strain, characterised by high virulence, was isolated from a latently infected American invasive signal crayfish (*Pacifastacus leniusculus*) host. The utilised Hap A strain, characterised by low virulence, was isolated from a repeatedly challenged, latently infected noble crayfish host population [138]. Consequently, both strains should represent extremes in the mosaic landscape of *A. astaci* strains present in Europe. The results of the infection experiment described in Francesconi et al. [33] showed that noble crayfish challenged with *A. astaci* Hap B have the highest amount of pathogen DNA inside their tissues, indicating that the pathogen successfully overcame the immune defences of the host. This corresponds to the high number of immune related DEGs observed in this experimental group. Furthermore, it was observed in other experiments that all the noble crayfish infected with this specific Hap B strain die within two weeks after challenge (our unpublished experimental results). Concurrently, Hap A challenged noble crayfish successfully contained the pathogen, without the apparent mobilisation of the hepatopancreas in the immune response and remained asymptomatic 45 days post-challenge [33]. However, Hap A challenged marbled crayfish showed the highest number of immune related DEGs to the non-associated pathogen strain, while in the Hap B challenged marbled crayfish no immune response activation was observed based on the differential gene expression analysis, with, however, the proPO, AMPs and recognition gene sets enriched, suggesting the possible involvement of these pathways (**Figure 5, Figure S2**). Contemporaneously, the highest amount of pathogen DNA in the marbled crayfish was detected in the Hap B challenged group [33]. This result indicates that the virulence of *A. astaci* and its ability to colonise the host’s tissues are not the only factors influencing the strength of the host’s immune response. In fact, one possible explanation could revolve around processes of coevolution between the crayfish and a specific strain of *A. astaci*.

It has been shown in several instances that invertebrates, although lacking an adaptive immune system, can build an immune memory, mounting an immune response of different magnitude after subsequent exposures to the same pathogen [18,139]. Such a response could be of tolerance with a lowered immune response to known stimuli, or of potentiation with a higher immune response upon re-encounter of the same pathogen [139]. Furthermore, transgenerational immune priming, in which the immune memory is transferred to the next generations by parents exposed to the pathogen, has been observed in insects [140,141] and in the brine shrimp (*Artemia franciscana*) [142]. While the specific mechanisms are not completely understood and are likely to be different depending on the host and the parasite, transgenerational immune priming might be the basis of the long-debated host-pathogen coevolution between North American crayfish and *A. Astaci* [8,143].

It is accepted that coevolution is a dynamic and ongoing process, in which the rapid adaptation of the host to the pathogen (and *vice versa*) can occur over short time frames, even a few decades [144]. The Hap A strain was isolated from latently infected noble crayfish in Lake Venesjärvi, Finland. The noble crayfish population in the lake faced at least 3 mass mortalities in the past 50 years until the year 2000. In 2013, the population was identified as carrier of *A. astaci* [138]. The results of our study suggest that, probably in the span of 50 years, the Hap A strain used in this study adapted to its naïve native European crayfish host, presumably through modification of its pathogenic epitopes. This has resulted in the overall lower virulence of the pathogen and lower immune stimulation of the host. At the same time, new epitopes presented by this *A. astaci* strain led to the higher expression of the diverse PRR genes in the marbled crayfish, responsible for the recognition of the pathogen and for boosting its immune response capability. However, considering that the gene expression analysis in marbled crayfish was conducted after removal of the batch effect related to reproducing crayfish, this could have biased our results. It has already been shown that immune related genes are over-expressed in reproducing insects [145]. If, similarly, reproduction in crayfish involves an up-regulation of immune related genes, the removal of the batch effect might have also removed relevant DEGs in the marble crayfish groups.

### 3.7. Study limitations

This study provides a deep insight into the innate immune response following an *A. astaci* challenge in the noble crayfish and the marbled crayfish. Transcriptomic data allowed us to explore the gene expression landscape and to identify key genes in the crayfish immunity. However, information about genomic locations and gene surroundings, which are highly influential on the gene expression profiles, are still not available. Consequently, generating first high-quality genome assemblies for freshwater crayfish represents a priority in the field of crayfish immunity, and would allow for the future comprehensive epigenomic studies. Unfortunately, until now this has proven to be a challenging task, because freshwater crayfish genomes are often large in size and have a high proportion of repetitive DNA sequences [31,146,147]. Furthermore, while in Decapods the role of the hepatopancreas in the immune response against pathogens has already been demonstrated, it has to be considered that the observed expression profile might be influenced by the infiltrating haemocytes [16,104]. In the future, this issue could be resolved by investigating additional tissues and by applying the higher resolution single cell RNA sequencing, capable of differentiating different cell populations within a tissue [148].

## 4. Conclusions

Identifying genes and pathways involved in the immune response to the pathogen *A. astaci* challenge represents a milestone in the conservation and aquaculture efforts for the native European crayfish species. Our analysis of the gene expression patterns in the noble crayfish and marbled crayfish highlighted a critical difference between the invasive and native species in response to the *A. astaci* challenge. Acutely infected noble crayfish in the response to Hap B strain of *A. astaci* (high virulence) relied mainly on the proPO cascade. The activation of this cascade results in the synthesis of highly toxic metabolites, capable of inflicting damage to the invading pathogen, but also to the host. On the other side, in the marbled crayfish infected with the Hap A strain of *A. astaci* (low virulence), we observed a mobilisation of PPRs (DSCAM, C-type lectins), AMPs (crustins and ALFs), and HCYs, capable of mimicking the proPO activity without inflicting damage to the host. A common denominator for both species was the absence of the clear late immune response (21 days post-challenge) once the immune tolerance to the invading pathogen was achieved. For the first time, we showcased the importance of the hepatopancreas as a highly relevant immune system organ in the response to the *A. astaci* challenge, for both the native noble crayfish and invasive marbled crayfish. The general overview of the freshwater crayfish immune response arsenal presented in this study will provide a backbone for the future advances in crayfish immunology.

## Abbreviations

DSCAM: down syndrome cell adhesion molecule
proPO: prophenoloxidase
PAMP: pathogen associated molecular pattern
PRP: pattern recognition proteins
TLR: toll-like receptor
Hap B: haplogroup B
Hap A: haplogroup A
DE: differentially expressed
DEG: differentially expressed gene
AMP: antimicrobial peptides
GNBPs: β-(1,3)-glucan receptors
ppA: serine protease
PXN: peroxinectin
PO: phenoloxidase
INX: innexin
HCY: hemocyanin
TPM: transcripts per million
Clip SP: serine proteinase
Serpin: serine proteinase inhibitor
MIP: melanisation inhibition protein
CPC-1-like: caspase 1-like molecule
CTL: C-type lectin
Ig: immunoglobulin, Fn, fibronectin
aa: amino acid
CRP: C-reactive protein
SAP: serum- amyloid P component
TWD: triple whey acidic protein
WAP: whey acidic protein
ALFs: antilipopolysaccharide factor
C/EBP: CCAAT/enhancer-binding protein
C/EBP-β: CCAAT/enhancer-binding protein beta
KLF1: krüppel 1-like factor protein
KLF: krüppel-like factor
Crus: crustin
TNFα: tumor necrosis factor alpha
TNF: tumour necrosis factor
AP: alkaline phosphatase
ETosis: extracellular trap release
HSP: heat-shock protein
WSSV: white spot syndrome virus.

## Author Contributions

K.T., C.F., J.J., J.M. **Conceptualization**; Lj.L.B., A.K., C.R. **Data curation**; Lj.L.B., C.F., C.R., L.H, L.P. **Formal analysis**; K.T., M.B. **Funding acquisition**; C.F., J.J., J.M., K.T. **Investigation**; Lj.L.B., O.L., C.R., L.H, L.P., B.F. **Methodology**; K.T. **Project administration**; K.T., O.L., M.B. **Resources**; A.K., Lj.L.B, C.R. **Software**; O.L., K.T., M.B. **Supervision**; O.L., K.T., C.F., Lj.L.B. **Validation**; Lj.L.B. **Visualization**; Lj.L.B., C.F. **Roles/Writing – original draft**; Lj.L.B., C.F., K.T., O.L., C.R., L.H., L.P., A.K., J.J., J.M., B.F., M.B. **Writing – review & editing**.

## Funding

This work was supported by the IdEx Unistra in the framework of the “Investments for the future” program of the French government and Institute funds from the Centre National de la Recherche Scientifique and the Université de Strasbourg.

K.T. and M.B. received seed funding for RNA sequencing from the LOEWE center for Translational Biodiversity Genomics (TBG).

## Acknowledgements

The authors would like to express their gratitude to Dr. Clement Schneider and Alexandra Schmidt for their helpful suggestions.

We would also like to acknowledge the support from Jorg Rapp in the server administration and the BIGEst platform for informatics support.

## Conflict of Interest Statement

The authors declare that they have no known competing financial interests or personal relationships which have or could be perceived to have influenced the work reported in this article.

## Supplementary files

**Table S1.** List of sequences used in the BLAST analysis for identification of the innate immunity genes in noble and marbled crayfish and their respective gene accession numbers.

**Table S2.** Innate immunity genes identified through the BLAST search with their respective match length, %identity, e-values and Dammit! annotations in the noble crayfish.

**Table S3.** Innate immunity genes identified through the BLAST search with their respective match length, %identity, e-values and Dammit! annotations in the marbled crayfish.

**Table S4.** Raw and post pre-processing Illumina sequence data statistics and mapping results of the read pseudo-alignment with Salmon against the *de novo* assembled transcriptome assemblies for noble and marbled crayfish.

**Table S5.** List of differentially expressed genes in the response of the noble crayfish to the challenge with *A. astaci*.

**Table S6.** List of differentially expressed genes and their respective annotations in the response of the marbled crayfish to the challenge with *A. astaci*.

**Figure S1.** Results of the principal component analysis (PCA) analysis for (a) noble crayfish and (b) marbled crayfish on the rlog transformed datasets, indicating batch effect related to differences between males (blue) and females (red) in noble crayfish and reproduction (reproducing-green, non-reproducing-purple) in marbled crayfish. The PCA with batch effect removal using removeBatchEffect() function implemented in limma R package (Ritchie et al., 2015) for (c) noble crayfish and (d) marbled crayfish.

**Figure S2**. Results of the Gene set enrichment analysis for (a) Hap A challenged noble crayfish (Day 3), (b) Hap B challenged noble crayfish (Day 21), (c) Hap B challenged marbled crayfish (Day 3), (d) Hap B challenged marbled crayfish (Day 21). Adjusted p-values, and Normalized enrichment scores (NES) are shown. AMPs- antimicrobial peptides, ProPO- prophenoloxidase pathway.

**File S1.** FASTA sequences used in the BLAST analysis for identification of the innate immunity genes in noble and marbled crayfish.

**File S2.** FASTA sequences of the innate immunity related transcripts identified through the BLAST analysis in the noble crayfish.

**File S3.** FASTA sequences of the innate immunity related transcripts identified through the BLAST analysis in the marbled crayfish.

## Notes

### Competing Interest Statement

The authors have declared no competing interest.

## References

[1] J. Reynolds, C. Souty-Grosset, A. Richardson, Ecological roles of crayfish in freshwater and terrestrial habitats, Freshw. Crayfish. 19 (2013) 197–218. https://doi.org/10.5869/fc.2013.v19-2.197.

[2] D.M. Holdich, J.D. Reynolds, C. Souty-Grosset, P.J. Sibley, A review of the ever increasing threat to European crayfish from non-indigenous crayfish species, Knowl. Manag. Aquat. Ecosyst. (2009) 11. https://doi.org/10.1051/kmae/2009025.

[3] D.J. Alderman, Geographical spread of bacterial and fungal diseases of crustaceans, Rev. Sci. Tech. l’OIE. 15 (1996) 603–632. https://doi.org/10.20506/rst.15.2.943.

[4] J. Makkonen, J. Jussila, J. Panteleit, N.S. Keller, A. Schrimpf, K. Theissinger, R. Kortet, L. Martín-Torrijos, J.V. Sandoval-Sierra, J. Diéguez-Uribeondo, H. Kokko, MtDNA allows the sensitive detection and haplotyping of the crayfish plague disease agent *Aphanomyces astaci* showing clues about its origin and migration, Parasitology. 145 (2018) 1210–1218. https://doi.org/10.1017/S0031182018000227.

[5] J. Makkonen, J. Jussila, R. Kortet, A. Vainikka, H. Kokko, Differing virulence of *Aphanomyces astaci* isolates and elevated resistance of noble crayfish *Astacus astacus* against crayfish plague, Dis. Aquat. Organ. 102 (2012) 129–136. https://doi.org/10.3354/dao02547.

[6] T. Becking, A. Mrugała, C. Delaunay, J. Svoboda, M. Raimond, S. Viljamaa-Dirks, A. Petrusek, F. Grandjean, C. Braquart-Varnier, Effect of experimental exposure to differently virulent Aphanomyces astaci strains on the immune response of the noble crayfish *Astacus astacus*, J. Invertebr. Pathol. 132 (2015) 115–124. https://doi.org/10.1016/j.jip.2015.08.007.

[7] L. Martín-Torrijos, M. Campos Llach, Q. Pou-Rovira, J. Diéguez-Uribeondo, Resistance to the crayfish plague, *Aphanomyces astaci* (Oomycota) in the endangered freshwater crayfish species, *Austropotamobius pallipes*, PLoS One. 12 (2017) 1–13. https://doi.org/10.1371/journal.pone.0181226.

[8] T. Unestam, Resistance to the crayfish plague in some American, Japanese and European crayfishes, Rep. Inst. Freshw. Res. Drottningholm. 49 (1969) 202–209.

[9] J. Jussila, A. Vrezec, J. Makkonen, R. Kortet, H. Kokko, Invasive crayfish and their invasive diseases in Europe with the focus on the virulence evolution of the crayfish plague, Biol. Invasions Chang. Ecosyst. Vectors, Ecol. Impacts, Manag. Predict. (2015) 183–211. https://doi.org/10.1515/9783110438666-013.

[10] L. Nyhlén, T. Unestam, Wound reactions and *Aphanomyces astaci* growth in crayfish cuticle, J. Invertebr. Pathol. 36 (1980) 187–197. https://doi.org/10.1016/0022-2011(80)90023-3.

[11] J. Jussila, A. Vrezec, T. Jaklič, H. Kukkonen, J. Makkonen, H. Kokko, *Aphanomyces astaci* isolate from latently infected stone crayfish (*Austropotamobius torrentium*) population is virulent, J. Invertebr. Pathol. 149 (2017) 15–20. https://doi.org/10.1016/j.jip.2017.07.003.

[12] J. Jussila, I. Maguire, H. Kokko, V. Tiitinen, J. Makkonen, Narrow-clawed crayfish in Finland: *Aphanomyces astaci* resistance and genetic relationship to other selected European and Asian populations, Knowl. Manag. Aquat. Ecosyst. 2020-Janua (2020). https://doi.org/10.1051/kmae/2020022.

[13] L. Cerenius, E. Bangyeekhun, P. Keyser, I. Söderhäll, K. Söderhäll, Host prophenoloxidase expression in freshwater crayfish is linked to increased resistance to the crayfish plague fungus, *Aphanomyces astaci*, Cell. Microbiol. 5 (2003) 353–357. https://doi.org/10.1046/j.1462-5822.2003.00282.x.

[14] C. Hauton, The scope of the crustacean immune system for disease control, J. Invertebr. Pathol. 110 (2012) 251–260. https://doi.org/10.1016/j.jip.2012.03.005.

[15] L. Cerenius, K. Söderhäll, Crayfish immunity – Recent findings, Dev. Comp. Immunol. 80 (2018) 94–98. https://doi.org/10.1016/j.dci.2017.05.010.

[16] A.F. Rowley, The Immune System of Crustaceans, Elsevier, 2016. https://doi.org/10.1016/B978-0-12-374279-7.12005-3.

[17] P. Jiravanichpaisal, B.L. Lee, K. Söderhäll, Cell-mediated immunity in arthropods: Hematopoiesis, coagulation, melanization and opsonization, Immunobiology. 211 (2006) 213–236. https://doi.org/10.1016/j.imbio.2005.10.015.

[18] C.F. Low, C.M. Chong, Peculiarities of innate immune memory in crustaceans, Fish Shellfish Immunol. 104 (2020) 605–612. https://doi.org/10.1016/j.fsi.2020.06.047.

[19] X. Lin, I. Söderhäll, Crustacean hematopoiesis and the astakine cytokines, Blood. 117 (2011) 6417–6424. https://doi.org/10.1182/blood-2010-11-320614.

[20] V.J. Smith, Immunology of Invertebrates: Cellular, ELS. (2016) 1–13. https://doi.org/10.1002/9780470015902.a0002344.pub3.

[21] P.T. Johnson, A review of fixed phagocytic and pinocytotic cells of decapod crustaceans, with remarks on hemocytes, Dev. Comp. Immunol. 11 (1987) 679–704. https://doi.org/10.1016/0145-305X(87)90057-7.

[22] T. Rőszer, The invertebrate midintestinal gland (“hepatopancreas”) is an evolutionary forerunner in the integration of immunity and metabolism, Cell Tissue Res. 358 (2014) 685–695. https://doi.org/10.1007/s00441-014-1985-7.

[23] V. Alday-Sanz, A. Roque, J. Turnbull, Clearing mechanisms of *Vibrio vulnificus* biotype I in the black tiger shrimp *Penaeus monodon*, Dis. Aquat. Organ. 48 (2002) 91–99. https://doi.org/10.3354/dao048091.

[24] D. Chen, L. Guo, C. Yi, S. Wang, Y. Ru, H. Wang, Ecotoxicology and Environmental Safety Hepatopancreatic transcriptome analysis and humoral immune factor assays in red claw crayfish (*Cherax quadricarinatus*) provide insight into innate immunomodulation under Vibrio parahaemolyticus infection, Ecotoxicol. Environ. Saf. 217 (2021) 112266. https://doi.org/10.1016/j.ecoenv.2021.112266.

[25] X. Meng, L. Hong, T.T. Yang, Y. Liu, T. Jiao, X.H. Chu, D.Z. Zhang, J.L. Wang, B.P. Tang, Q.N. Liu, W.W. Zhang, W.F. He, Transcriptome-wide identification of differentially expressed genes in *Procambarus clarkii* in response to chromium challenge, Fish Shellfish Immunol. 87 (2019) 43–50. https://doi.org/10.1016/j.fsi.2018.12.055.

[26] L.S. Dai, M.N. Abbas, S. Kausar, Y. Zhou, Transcriptome analysis of hepatopancraes of *Procambarus clarkii* challenged with polyriboinosinic polyribocytidylic acid (poly I:C), Fish Shellfish Immunol. 71 (2017) 144–150. https://doi.org/10.1016/j.fsi.2017.10.010.

[27] T. Jiao, T.T. Yang, D. Wang, Z.Q. Gao, J.L. Wang, B.P. Tang, Q.N. Liu, D.Z. Zhang, L.S. Dai, Characterization and expression analysis of immune-related genes in the red swamp crayfish, *Procambarus clarkii* in response to lipopolysaccharide challenge, Fish Shellfish Immunol. 95 (2019) 140–150. https://doi.org/10.1016/j.fsi.2019.09.072.

[28] G. Shen, X. Zhang, J. Gong, Y. Wang, P. Huang, Y. Shui, Z. Xu, H. Shen, Transcriptomic analysis of *Procambarus clarkii* affected by “Black May” disease, Sci. Rep. 10 (2020) 1–13. https://doi.org/10.1038/s41598-020-78191-8.

[29] Y. Zhang, Z. Li, S. Kholodkevich, A. Sharov, Y. Feng, N. Ren, K. Sun, Cadmium-induced oxidative stress, histopathology, and transcriptome changes in the hepatopancreas of freshwater crayfish (*Procambarus clarkii*), Sci. Total Environ. 666 (2019) 944–955. https://doi.org/10.1016/j.scitotenv.2019.02.159.

[30] G. Vogt, Investigating the genetic and epigenetic basis of big biological questions with the parthenogenetic marbled crayfish: A review and perspectives, J. Biosci. 43 (2018) 189–223. https://doi.org/10.1007/s12038-018-9741-x.

[31] J. Gutekunst, R. Andriantsoa, C. Falckenhayn, K. Hanna, W. Stein, J. Rasamy, F. Lyko, Clonal genome evolution and rapid invasive spread of the marbled crayfish, Nat. Ecol. Evol. 2 (2018) 567–573. https://doi.org/10.1038/s41559-018-0467-9.

[32] N.S. Keller, M. Pfeiffer, I. Roessink, R. Schulz, A. Schrimpf, First evidence of crayfish plague agent in populations of the marbled crayfish (*Procambarus fallax* forma virginalis), Knowl. Manag. Aquat. Ecosyst. (2014) 15. https://doi.org/10.1051/kmae/2014032.

[33] C. Francesconi, J. Makkonen, A. Schrimpf, J. Jussila, H. Kokko, K. Theissinger, Controlled infection experiment with *Aphanomyces astaci* provides evidence for latent infections and resistance in freshwater crayfish, Accept. Manuscr. (n.d.).

[34] L.L. Boštjančić, C. Francesconi, C. Rutz, L. Hoffbeck, L. Poidevin, A. Kress, J. Jussila, J. Makkonen, B. Feldmeyer, M. Bálint, O. Lecompte, K. Theissinger, Dataset of the *de novo* assembly and annotation of the marbled crayfish and noble crayfish hepatopancreas transcriptomes, Submitt. Manuscr. (n.d.).

[35] A.G. Lai, A.A. Aboobaker, Comparative genomic analysis of innate immunity reveals novel and conserved components in crustacean food crop species, BMC Genomics. 18 (2017) 1–26. https://doi.org/10.1186/s12864-017-3769-4.

[36] B. Buchfink, C. Xie, D.H. Huson, Fast and sensitive protein alignment using DIAMOND, Nat. Methods. 12 (2015) 59–60. https://doi.org/10.1038/nmeth.3176.

[37] B.D. Ondov, N.H. Bergman, A.M. Phillippy, Interactive metagenomic visualization in a Web browser, BMC Bioinformatics. 12 (2011) 385. https://doi.org/10.1186/1471-2105-12-385.

[38] R. Patro, G. Duggal, M.I. Love, R.A. Irizarry, C. Kingsford, Salmon provides fast and bias-aware quantification of transcript expression, Nat. Methods. 14 (2017) 417–419. https://doi.org/10.1038/nmeth.4197.

[39] A. Srivastava, L. Malik, H. Sarkar, M. Zakeri, F. Almodaresi, C. Soneson, M.I. Love, C. Kingsford, R. Patro, Alignment and mapping methodology influence transcript abundance estimation, Genome Biol. 21 (2020) 1–21. https://doi.org/10.1186/s13059-020-02151-8.

[40] A. Roberts, C. Trapnell, J. Donaghey, J.L. Rinn, L. Pachter, Improving RNA-Seq expression estimates by correcting for fragment bias, Genome Biol. 12 (2011) R22. https://doi.org/10.1186/gb-2011-123-r22.

[41] M.I. Love, W. Huber, S. Anders, Moderated estimation of fold change and dispersion for RNA-seq data with DESeq2, Genome Biol. 15 (2014) 1–21. https://doi.org/10.1186/s13059-014-0550-8.

[42] C. Soneson, M.I. Love, M.D. Robinson, Differential analyses for RNA-seq: transcript-level estimates improve gene-level inferences, F1000Research. 4 (2015) 1521. https://doi.org/10.12688/f1000research.7563.1.

[43] K. Blighe, S. Rana, M. Lewis, EnhancedVolcano: Publication-ready volcano plots with enhanced colouring and labeling, Available at: https://Github.Com/Kevinblighe/EnhancedVolcano. (2020).

[44] A. Zhu, J.G. Ibrahim, M.I. Love, Heavy-tailed prior distributions for sequence count data: removing the noise and preserving large differences, Bioinformatics. 35 (2019) 2084–2092. https://doi.org/10.1093/bioinformatics/bty895.

[45] M.E. Ritchie, B. Phipson, D. Wu, Y. Hu, C.W. Law, W. Shi, G.K. Smyth, limma powers differential expression analyses for RNA-sequencing and microarray studies, Nucleic Acids Res. 43 (2015) e47–e47. https://doi.org/10.1093/nar/gkv007.

[46] G. Yu, L.-G. Wang, Y. Han, Q.-Y. He, clusterProfiler: an R Package for Comparing Biological Themes Among Gene Clusters, Omi. A J. Integr. Biol. 16 (2012) 284–287. https://doi.org/10.1089/omi.2011.0118.

[47] K.F. Clark, S.J. Greenwood, Next-generation sequencing and the crustacean immune system: The need for alternatives in immune gene annotation, Integr. Comp. Biol. 56 (2016) 1113–1130. https://doi.org/10.1093/icb/icw023.

[48] G. Calderón-Rosete, J.A. González-Barrios, M. Lara-Lozano, C. Piña-Leyva, L. Rodríguez-Sosa, Transcriptional identification of related proteins in the immune system of the crayfish *Procambarus clarkii*, High-Throughput. 7 (2018) 1–15. https://doi.org/10.3390/HT7030026.

[49] L. Cerenius, K. Söderhäll, The prophenoloxidase-activating system in invertebrates, Immunol. Rev. 198 (2004) 116–126. https://doi.org/10.1111/j.0105-2896.2004.00116.x.

[50] S. Paro, J.-L. Imler, Immunity in insects, Encycl. Immunol. Elsevier Sci. 68 (2016) 383–398.

[51] T. Kawasaki, T. Kawai, Toll-like receptor signaling pathways, Front. Immunol. 5 (2014) 1–8. https://doi.org/10.3389/fimmu.2014.00461.

[52] C.A. Janeway, R. Medzhitov, Innate immune recognition, Annu. Rev. Immunol. 20 (2002) 197–216. https://doi.org/10.1146/annurev.immunol.20.083001.084359.

[53] L. Cerenius, K. Söderhäll, Crustacean immune responses and their implications for disease control, in: Infect. Dis. Aquac., Elsevier, 2012: pp. 69–87. https://doi.org/10.1533/9780857095732.1.69.

[54] S. Shakibazadeh, C.R. Saad, A. Christianus, M.S. Kamarudin, K. Sijam, M. Nor Shamsudin, V.K. Neela, Bacteria flora associated with different body parts of hatchery reared juvenile *Penaeus monodon*, tanks water and sediment, Ann. Microbiol. 59 (2009) 425–430. https://doi.org/10.1007/BF03175126.

[55] M.K. Cheung, H.Y. Yip, W. Nong, P.T.W. Law, K.H. Chu, H.S. Kwan, J.H.L. Hui, Rapid Change of Microbiota Diversity in the Gut but Not the Hepatopancreas During Gonadal Development of the New Shrimp Model *Neocaridina denticulata*, Mar. Biotechnol. 17 (2015) 811–819. https://doi.org/10.1007/s10126-015-9662-8.

[56] F. Cornejo-Granados, A.A. Lopez-Zavala, L. Gallardo-Becerra, A. Mendoza-Vargas, F. Sánchez, R. Vichido, L.G. Brieba, M.T. Viana, R.R. Sotelo-Mundo, A. Ochoa-Leyva, Microbiome of Pacific Whiteleg shrimp reveals differential bacterial community composition between Wild, Aquacultured and AHPND/EMS outbreak conditions, Sci. Rep. 7 (2017) 1–15. https://doi.org/10.1038/s41598-017-11805-w.

[57] K. Orlić, L. Šver, L. Burić, S. Kazazić, D. Grbin, I. Maguire, D. Pavić, R. Hrašćan, T. Vladušić, S. Hudina, A. Bielen, Cuticle-associated bacteria can inhibit crayfish pathogen *Aphanomyces astaci*: Opening the perspective of biocontrol in astaciculture, Aquaculture. 533 (2021) 736112. https://doi.org/10.1016/j.aquaculture.2020.736112.

[58] M.C. Ooi, E.F. Goulden, G.G. Smith, A.R. Bridle, Haemolymph microbiome of the cultured spiny lobster *Panulirus ornatus* at different temperatures, Sci. Rep. 9 (2019) 1–13. https://doi.org/10.1038/s41598-019-39149-7.

[59] X.-W. Wang, J.-X. Wang, Crustacean hemolymph microbiota: Endemic, tightly controlled, and utilization expectable, Mol. Immunol. 68 (2015) 404–411. https://doi.org/10.1016/j.molimm.2015.06.018.

[60] L. Cerenius, B.L. Lee, K. Söderhäll, The proPO-system: pros and cons for its role in invertebrate immunity, Trends Immunol. 29 (2008) 263–271. https://doi.org/10.1016/j.it.2008.02.009.

[61] A. Subramanian, P. Tamayo, V.K. Mootha, S. Mukherjee, B.L. Ebert, M.A. Gillette, A. Paulovich, S.L. Pomeroy, T.R. Golub, E.S. Lander, J.P. Mesirov, Gene set enrichment analysis: A knowledge-based approach for interpreting genome-wide expression profiles, Proc. Natl. Acad. Sci. U. S. A. 102 (2005) 15545–15550. https://doi.org/10.1073/pnas.0506580102.

[62] K. Söderhäll, L. Cerenius, Role of the prophenoloxidase-activating system in invertebrate immunity, Curr. Opin. Immunol. 10 (1998) 23–28. https://doi.org/10.1016/S0952-7915(98)80026-5.

[63] M. Jearaphunt, C. Noonin, P. Jiravanichpaisal, S. Nakamura, A. Tassanakajon, I. Söderhäll, K. Söderhäll, Caspase-1-Like Regulation of the proPO-System and Role of ppA and Caspase-1-Like Cleaved Peptides from proPO in Innate Immunity, PLoS Pathog. 10 (2014). https://doi.org/10.1371/journal.ppat.1004059.

[64] M.W. Johansson, T. Holmblad, P.O. Thörnqvist, M. Cammarata, N. Parrinello, K. Söderhäll, A cell-surface superoxide dismutase is a binding protein for peroxinectin, a cell-adhesive peroxidase in crayfish., J. Cell Sci. 112 (Pt 6 (1999) 917–25. http://www.ncbi.nlm.nih.gov/pubmed/10036241.

[65] X. Lin, L. Cerenius, B.L. Lee, K. Söderhäll, Purification of properoxinectin, a myeloperoxidase homologue and its activation to a cell adhesion molecule, Biochim. Biophys. Acta – Gen. Subj. 1770 (2007) 87–93. https://doi.org/10.1016/j.bbagen.2006.06.018.

[66] L. Cerenius, K. Söderhäll, Arthropoda: Pattern Recognition Proteins in Crustacean Immunity, in: Adv. Comp. Immunol., Springer International Publishing, Cham, 2018: pp. 213–224. https://doi.org/10.1007/978-3-319-76768-0_10.

[67] Z. Liang, L. Sottrup-Jensen, A. Aspan, M. Hall, K. Soderhall, Pacifastin, a novel 155- kDa heterodimeric proteinase inhibitor containing a unique transferrin chain, Proc. Natl. Acad. Sci. 94 (1997) 6682–6687. https://doi.org/10.1073/pnas.94.13.6682.

[68] I. Söderhäll, C. Wu, M. Novotny, B.L. Lee, K. Söderhäll, A novel protein acts as a negative regulator of prophenoloxidase activation and melanization in the freshwater crayfish *Pacifastacus leniusculus*, J. Biol. Chem. 284 (2009) 6301–6310. https://doi.org/10.1074/jbc.M806764200.

[69] E. De Gregorio, S.J. Han, W.J. Lee, M.J. Baek, T. Osaki, S. Kawabata, B.L. Lee, S. Iwanaga, B. Lemaitre, P.T. Brey, An, immune-responsive Serpin regulates the melanization cascade, Dros. Dev. Cell. 3 (2002) 581–592.

[70] E. Bangyeekhun, L. Cerenius, K. Söderhäll, Molecular cloning and characterization of two serine proteinase genes from the crayfish plague fungus, *Aphanomyces astaci*, J. Invertebr. Pathol. 77 (2001) 206–216. https://doi.org/10.1006/jipa.2001.5019.

[71] Y.-R. Zhao, Y.-H. Xu, H.-S. Jiang, S. Xu, X.-F. Zhao, J.-X. Wang, Antibacterial activity of serine protease inhibitor 1 from kuruma shrimp Marsupenaeus japonicus, Dev. Comp. Immunol. 44 (2014) 261–269. https://doi.org/10.1016/j.dci.2014.01.002.

[72] T. Burmester, Origin and evolution of arthropod hemocyanins and related proteins, J. Comp. Physiol. B Biochem. Syst. Environ. Physiol. 172 (2002) 95–107. https://doi.org/10.1007/s00360-001-0247-7.

[73] H. Decker, N. Hellmann, E. Jaenicke, B. Lieb, U. Meissner, J. Markl, Minireview: Recent progress in hemocyanin research, Integr. Comp. Biol. 47 (2007) 631–644. https://doi.org/10.1093/icb/icm063.

[74] S.Y. Lee, B.L. Lee, K. Söderhäll, Processing of crayfish hemocyanin subunits into phenoloxidase, Biochem. Biophys. Res. Commun. 322 (2004) 490–496. https://doi.org/10.1016/j.bbrc.2004.07.145.

[75] D.A. Ward, E.M. Sefton, M.C. Prescott, S.G. Webster, G. Wainwright, H.H. Rees, M.J. Fisher, Efficient identification of proteins from ovaries and hepatopancreas of the unsequenced edible crab, *Cancer pagurus*, by mass spectrometry and homology-based, cross-species searching, J. Proteomics. 73 (2010) 2354–2364. https://doi.org/10.1016/j.jprot.2010.07.008.

[76] N.A. Ratcliffe, A.F. Rowley, S.W. Fitzgerald, C.P. Rhodes, Invertebrate Immunity: Basic Concepts and Recent Advances, in: 1985: pp. 183–350. https://doi.org/10.1016/S0074-7696(08)62351-7.

[77] R., Paul, B., Bergner, A., Pfeffer-Seidl, H., Decker, R., Efinger, H., Storz, Gas transport in the haemolymph of arachnids – oxygen transport and the physiological role of haemocyanin, J. Exp. Biol. 188 (1994) 25–46. http://www.ncbi.nlm.nih.gov/pubmed/9317270.

[78] H. Choi, D.G. Lee, Antifungal activity and pore-forming mechanism of astacidin 1 against *Candida albicans*, Biochimie. 105 (2014) 58–63. https://doi.org/10.1016/j.biochi.2014.06.014.

[79] H. Lis, N. Sharon, Lectins: Carbohydrate-Specific Proteins That Mediate Cellular Recognition, Chem. Rev. 98 (1998) 637–674. https://doi.org/10.1021/cr940413g.

[80] X.-K. Jin, S. Li, X.-N. Guo, L. Cheng, M.-H. Wu, S.-J. Tan, Y.-T. Zhu, A.-Q. Yu, W.-W. Li, Q. Wang, Two antibacterial C-type lectins from crustacean, *Eriocheir sinensis*, stimulated cellular encapsulation in vitro, Dev. Comp. Immunol. 41 (2013) 544–552. https://doi.org/10.1016/j.dci.2013.07.016.

[81] X.-Z. Shi, L. Wang, S. Xu, X.-W. Zhang, X.-F. Zhao, G.R. Vasta, J.-X. Wang, A Galectin from the Kuruma Shrimp (*Marsupenaeus japonicus*) Functions as an Opsonin and Promotes Bacterial Clearance from Hemolymph, PLoS One. 9 (2014) e91794. https://doi.org/10.1371/journal.pone.0091794.

[82] E. Ling, X. Yu, Cellular encapsulation and melanization are enhanced by immulectins, pattern recognition receptors from the tobacco hornworm *Manduca sexta*, Dev. Comp. Immunol. 30 (2006) 289–299. https://doi.org/10.1016/j.dci.2005.05.005.

[83] G.R. Vasta, Roles of galectins in infection, Nat. Rev. Microbiol. 7 (2009) 424–438. https://doi.org/10.1038/nrmicro2146.

[84] Z.-Y. Zhao, Z.-X. Yin, X.-P. Xu, S.-P. Weng, X.-Y. Rao, Z.-X. Dai, Y.-W. Luo, G. Yang, Z.-S. Li, H.-J. Guan, S.-D. Li, S.-M. Chan, X.-Q. Yu, J.-G. He, A Novel C-Type Lectin from the Shrimp *Litopenaeus vannamei* Possesses Anti-White Spot Syndrome Virus Activity, J. Virol. 83 (2009) 347–356. https://doi.org/10.1128/JVI.00707-08.

[85] L. Cerenius, S. ichiro Kawabata, B.L. Lee, M. Nonaka, K. Söderhäll, Proteolytic cascades and their involvement in invertebrate immunity, Trends Biochem. Sci. 35 (2010) 575–583. https://doi.org/10.1016/j.tibs.2010.04.006.

[86] K. Yamakawa, DSCAM: a novel member of the immunoglobulin superfamily maps in a Down syndrome region and is involved in the development of the nervous system, Hum. Mol. Genet. 7 (1998) 227–237. https://doi.org/10.1093/hmg/7.2.227.

[87] T.H. Ng, Y.A. Chiang, Y.C. Yeh, H.C. Wang, Review of Dscam-mediated immunity in shrimp and other arthropods, Dev. Comp. Immunol. 46 (2014) 129–138. https://doi.org/10.1016/j.dci.2014.04.002.

[88] T.H. Ng, R. Kumar, K. Apitanyasai, S.T. He, S.P. Chiu, H.C. Wang, Selective expression of a “correct cloud” of Dscam in crayfish survivors after second exposure to the same pathogen, Fish Shellfish Immunol. 92 (2019) 430–437. https://doi.org/10.1016/j.fsi.2019.06.023.

[89] P.-H. Chou, H.-S. Chang, I.-T. Chen, C.-W. Lee, H.-Y. Hung, K.C. Han-Ching Wang, *Penaeus monodon* Dscam (PmDscam) has a highly diverse cytoplasmic tail and is the first membrane-bound shrimp Dscam to be reported, Fish Shellfish Immunol. 30 (2011) 1109–1123. https://doi.org/10.1016/j.fsi.2011.02.009.

[90] B. Pees, W. Yang, A. Zárate-Potes, H. Schulenburg, K. Dierking, High Innate Immune Specificity through Diversified C-Type Lectin-Like Domain Proteins in Invertebrates, J. Innate Immun. 8 (2016) 129–142. https://doi.org/10.1159/000441475.

[91] E.C. Liebl, Interactions between the secreted protein Amalgam, its transmembrane receptor Neurotactin and the Abelson tyrosine kinase affect axon pathfinding, Development. 130 (2003) 3217–3226. https://doi.org/10.1242/dev.00545.

[92] T. Zeev-Ben-Mordehai, E. Mylonas, A. Paz, Y. Peleg, L. Toker, I. Silman, D.I. Svergun, J.L. Sussman, The Quaternary Structure of Amalgam, a Drosophila Neuronal Adhesion Protein, Explains Its Dual Adhesion Properties, Biophys. J. 97 (2009) 2316–2326. https://doi.org/10.1016/j.bpj.2009.07.045.

[93] T.W. Du Clos, Pentraxins: Structure, Function, and Role in Inflammation, ISRN Inflamm. 2013 (2013) 1–22. https://doi.org/10.1155/2013/379040.

[94] A. Mantovani, C. Garlanda, A. Doni, B. Bottazzi, Pentraxins in innate immunity: From C-reactive protein to the long pentraxin PTX3, J. Clin. Immunol. 28 (2008) 1–13. https://doi.org/10.1007/s10875-007-9126-7.

[95] P.B. Armstrong, Comparative Biology of the Pentraxin Protein Family: Evolutionarily Conserved Component of Innate Immune System, Elsevier Ltd, 2015. https://doi.org/10.1016/bs.ircmb.2015.01.002.

[96] Y.J. Ma, P. Garred, Pentraxins in Complement Activation and Regulation, Front. Immunol. 9 (2018) 3046. https://doi.org/10.3389/fimmu.2018.03046.

[97] M.B. Pepys, G.M. Hirschfield, C-reactive protein: a critical update, J. Clin. Invest. 111 (2003) 1805–1812. https://doi.org/10.1172/JCI18921.

[98] P.M.L. Ng, Z. Jin, S.S.H. Tan, B. Ho, J.L. Ding, C-reactive protein: a predominant LPS-binding acute phase protein responsive to *Pseudomonas infection*, J. Endotoxin Res. 10 (2004) 163–174. https://doi.org/10.1179/096805104225004833.

[99] K.F. Clark, A.R. Acorn, S.J. Greenwood, Differential expression of American lobster (*Homarus americanus*) immune related genes during infection of *Aerococcus viridans* var. homari, the causative agent of Gaffkemia, J. Invertebr. Pathol. 112 (2013) 192–202. https://doi.org/10.1016/j.jip.2012.11.005.

[100] K.F. Clark, A.R. Acorn, S.J. Greenwood, A transcriptomic analysis of American lobster (*Homarus americanus*) immune response during infection with the bumper car parasite *Anophryoides haemophila*, Dev. Comp. Immunol. 40 (2013) 112–122. https://doi.org/10.1016/j.dci.2013.02.009.

[101] A. Tassanakajon, K. Somboonwiwat, P. Amparyup, Sequence diversity and evolution of antimicrobial peptides in invertebrates, Dev. Comp. Immunol. 48 (2015) 324–341. https://doi.org/10.1016/j.dci.2014.05.020.

[102] V.J. Smith, E.A. Dyrynda, Antimicrobial proteins: From old proteins, new tricks, Mol. Immunol. 68 (2015) 383–398. https://doi.org/10.1016/j.molimm.2015.08.009.

[103] S. Sricharoen, J.J. Kim, S. Tunkijjanukij, I. Söderhäll, Exocytosis and proteomic analysis of the vesicle content of granular hemocytes from a crayfish, Dev. Comp. Immunol. 29 (2005) 1017–1031. https://doi.org/10.1016/j.dci.2005.03.010.

[104] Y.-P. Jia, Y.-D. Sun, Z.-H. Wang, Q. Wang, X.-W. Wang, X.-F. Zhao, J.-X. Wang, A single whey acidic protein domain (SWD)-containing peptide from fleshy prawn with antimicrobial and proteinase inhibitory activities, Aquaculture. 284 (2008) 246–259. https://doi.org/10.1016/j.aquaculture.2008.07.046.

[105] Y.X. Zhang, J.X. Wang, X.W. Wang, First identification and characterization of a triple WAP domain containing protein in *Procambarus clarkii* provides new insights into the classification and evolution of WAP proteins in crustacean, Fish Shellfish Immunol. 94 (2019) 592–598. https://doi.org/10.1016/j.fsi.2019.09.023.

[106] T. Becking, C. Delaunay, R. Cordaux, J.M. Berjeaud, C. Braquart-Varnier, J. Verdon, Shedding light on the antimicrobial peptide arsenal of terrestrial isopods: Focus on armadillidins, a new crustacean AMP family, Genes (Basel). 11 (2020). https://doi.org/10.3390/genes11010093.

[107] C. Sun, W.T. Xu, H.W. Zhang, L.P. Dong, T. Zhang, X.F. Zhao, J.X. Wang, An anti-lipopolysaccharide factor from red swamp crayfish, *Procambarus clarkii*, exhibited antimicrobial activities in vitro and in vivo, Fish Shellfish Immunol. 30 (2011) 295–303. https://doi.org/10.1016/j.fsi.2010.10.022.

[108] E. de la Vega, N.A. O’Leary, J.E. Shockey, J. Robalino, C. Payne, C.L. Browdy, G.W. Warr, P.S. Gross, Anti-lipopolysaccharide factor in *Litopenaeus vannamei* (LvALF): A broad spectrum antimicrobial peptide essential for shrimp immunity against bacterial and fungal infection, Mol. Immunol. 45 (2008) 1916–1925. https://doi.org/10.1016/j.molimm.2007.10.039.

[109] R. Bauer, C. Lehmann, J. Martini, F. Eckardt, M. Hoch, Gap Junction Channel Protein Innexin 2 Is Essential for Epithelial Morphogenesis in the Drosophila Embryo, Mol. Biol. Cell. 15 (2004) 2992–3004. https://doi.org/10.1091/mbc.e04-01-0056.

[110] J.C. Sáez, M.C. Brañes, L.A. Corvalán, E.A. Eugenin, H. González, A.D. Martínez, F. Palisson, Gap junctions in cells of the immune system: Structure, regulation and possible functional roles, Brazilian J. Med. Biol. Res. 33 (2000) 447–455. https://doi.org/10.1590/S0100-879X2000000400011.

[111] J. Neijssen, B. Pang, J. Neefjes, Gap junction-mediated intercellular communication in the immune system, Prog. Biophys. Mol. Biol. 94 (2007) 207–218. https://doi.org/10.1016/j.pbiomolbio.2007.03.008.

[112] E.J. Furshpan, D.D. Potter, Transmission at the giant motor synapses of the crayfish, J. Physiol. 145 (1959) 289–325. https://doi.org/10.1113/jphysiol.1959.sp006143.

[113] M.D. Adams, The Genome Sequence of *Drosophila melanogaster*, Science (80-.). 287 (2000) 2185–2195. https://doi.org/10.1126/science.287.5461.2185.

[114] B. Kandarian, J. Sethi, A. Wu, M. Baker, N. Yazdani, E. Kym, A. Sanchez, L. Edsall, T. Gaasterland, E. Macagno, The medicinal leech genome encodes 21 innexin genes: different combinations are expressed by identified central neurons, Dev. Genes Evol. 222 (2012) 29–44. https://doi.org/10.1007/s00427-011-0387-z.

[115] T. Starich, M. Sheehan, J. Jadrich, J. Shaw, Innexins in C. elegans, Cell Commun. Adhes. 8 (2001) 311–314. https://doi.org/10.3109/15419060109080744.

[116] S. Shruti, D.J. Schulz, K.M. Lett, E. Marder, Electrical coupling and innexin expression in the stomatogastric ganglion of the crab *Cancer borealis*, J. Neurophysiol. 112 (2014) 2946–2958. https://doi.org/10.1152/jn.00536.2014.

[117] S.P. Wang, F.Y. Chen, L.X. Dong, Y.Q. Zhang, H.Y. Chen, K. Qiao, K.J. Wang, A novel innexin2 forming membrane hemichannel exhibits immune responses and cell apoptosis in *Scylla paramamosain*, Fish Shellfish Immunol. 47 (2015) 485–499. https://doi.org/10.1016/j.fsi.2015.09.028.

[118] J. Güiza, I. Barría, J.C. Sáez, J.L. Vega, Innexins: Expression, regulation, and functions, Front. Physiol. 9 (2018) 1–9. https://doi.org/10.3389/fphys.2018.01414.

[119] D.P. Ramji, P. Foka, CCAAT/enhancer-binding proteins: Structure, function and regulation, Biochem. J. 365 (2002) 561–575. https://doi.org/10.1042/BJ20020508.

[120] W. Wang, X. Xia, L. Mao, S. Wang, The CCAAT/Enhancer-Binding Protein Family: Its Roles in MDSC Expansion and Function, Front. Immunol. 10 (2019) 1804. https://doi.org/10.3389/fimmu.2019.01804.

[121] P. Kamsaeng, A. Tassanakajon, K. Somboonwiwat, Regulation of antilipopolysaccharide factors, ALFPm3 and ALFPm6, in Penaeus monodon, Sci. Rep. 7 (2017) 1–13. https://doi.org/10.1038/s41598-017-12137-5.

[122] P. Amparyup, H. Kondo, I. Hirono, T. Aoki, A. Tassanakajon, Molecular cloning, genomic organization and recombinant expression of a crustin-like antimicrobial peptide from black tiger shrimp *Penaeus monodon*, Mol. Immunol. 45 (2008) 1085–1093. https://doi.org/10.1016/j.molimm.2007.07.031.

[123] J. Tsukada, Y. Yoshida, Y. Kominato, P.E. Auron, The CCAAT/enhancer (C/EBP) family of basic-leucine zipper (bZIP) transcription factors is a multifaceted highly-regulated system for gene regulation, Cytokine. 54 (2011) 6–19. https://doi.org/10.1016/j.cyto.2010.12.019.

[124] N.M. Pollak, M. Hoffman, I.J. Goldberg, K. Drosatos, Krüppel-Like Factors: Crippling and Uncrippling Metabolic Pathways, JACC Basic to Transl. Sci. 3 (2018) 132–156. https://doi.org/10.1016/j.jacbts.2017.09.001.

[125] K. Czakai, I. Leonhardt, A. Dix, M. Bonin, J. Linde, H. Einsele, O. Kurzai, J. Loeffler, Krüppel-like Factor 4 modulates interleukin-6 release in human dendritic cells after in vitro stimulation with Aspergillus fumigatus and *Candida albicans*, Sci. Rep. 6 (2016) 1–9. https://doi.org/10.1038/srep27990.

[126] P.H. Huang, S.C. Lu, S.H. Yang, P.S. Cai, C.F. Lo, L.K. Chang, Regulation of the immediate-early genes of white spot syndrome virus by *Litopenaeus vannamei* kruppel-like factor (LvKLF), Dev. Comp. Immunol. 46 (2014) 364–372. https://doi.org/10.1016/j.dci.2014.05.012.

[127] W.J. Liu, C.F. Lo, G.H. Kou, J.H. Leu, Y.J. Lai, L.K. Chang, Y.S. Chang, The promoter of the white spot syndrome virus immediate-early gene WSSV108 is activated by the cellular KLF transcription factor, Dev. Comp. Immunol. 49 (2015) 7–18. https://doi.org/10.1016/j.dci.2014.10.015.

[128] Y. Huang, Q. Ren, A Kruppel-like factor from *Macrobrachium rosenbergii* (MrKLF) involved in innate immunity against pathogen infection, Fish Shellfish Immunol. 95 (2019) 519–527. https://doi.org/10.1016/j.fsi.2019.10.070.

[129] M. Kim, J.H. Lee, S.Y. Lee, E. Kim, J. Chung, Caspar, a suppressor of antibacterial immunity in Drosophila, Proc. Natl. Acad. Sci. 103 (2006) 16358–16363. https://doi.org/10.1073/pnas.0603238103.

[130] F.R. Balkwill, Cytokines, in: Encycl. Life Sci., John Wiley & Sons, Ltd, Chichester, UK, 2001. https://doi.org/10.1038/npg.els.0000929.

[131] A.B. Guimarães-Costa, M.T.C. Nascimento, A.B. Wardini, L.H. Pinto-Da-Silva, E.M. Saraiva, ETosis: A microbicidal mechanism beyond cell death, J. Parasitol. Res. 2012 (2012). https://doi.org/10.1155/2012/929743.

[132] G. Bidla, M.S. Dushay, U. Theopold, Crystal cell rupture after injury in Drosophila requires the JNK pathway, small GTPases and the TNF homolog eiger, J. Cell Sci. 120 (2007) 1209–1215. https://doi.org/10.1242/jcs.03420.

[133] A. Kulkarni, J.H.W.M. Rombout, I.S.B. Singh, N.S. Sudheer, J.M. Vlak, C.M.A. Caipang, M.F. Brinchmann, V. Kiron, Truncated VP28 as oral vaccine candidate against WSSV infection in shrimp: An uptake and processing study in the midgut of *Penaeus monodon*, Fish Shellfish Immunol. 34 (2013) 159–166. https://doi.org/10.1016/j.fsi.2012.10.028.

[134] R.M. Vabulas, S. Raychaudhuri, M. Hayer-Hartl, F.U. Hartl, Protein folding in the cytoplasm and the heat shock response., Cold Spring Harb. Perspect. Biol. 2 (2010). https://doi.org/10.1101/cshperspect.a004390.

[135] H. Xu, F. Yan, X. Deng, J. Wang, T. Zou, X. Ma, X. Zhang, Y. Qi, The interaction of white spot syndrome virus envelope protein VP28 with shrimp Hsc70 is specific and ATP-dependent, Fish Shellfish Immunol. 26 (2009) 414–421. https://doi.org/10.1016/j.fsi.2009.01.001.

[136] Y. Zeng, C.-P. Lu, Identification of differentially expressed genes in haemocytes of the crayfish (*Procambarus clarkii*) infected with white spot syndrome virus by suppression subtractive hybridization and cDNA microarrays, Fish Shellfish Immunol. 26 (2009) 646–650. https://doi.org/10.1016/j.fsi.2008.11.005.

[137] R. Arya, M. Mallik, S.C. Lakhotia, Heat shock genes — integrating cell survival and death, J. Biosci. 32 (2007) 595–610. https://doi.org/10.1007/s12038-007-0059-3.

[138] J. Jussila, C. Francesconi, K. Theissinger, H. Kokko, J. Makkonen, Is *Aphanomyces astaci* losing its stamina: a latent crayfish plague disease agent from lake Venesjärvi, Finland, Submitt. Manuscr. (n.d.).

[139] D. Melillo, R. Marino, P. Italiani, D. Boraschi, Innate Immune Memory in Invertebrate Metazoans: A Critical Appraisal, Front. Immunol. 9 (2018) 1915. https://doi.org/10.3389/fimmu.2018.01915.

[140] S.M. Barribeau, P. Schmid-Hempel, B.M. Sadd, Royal Decree: Gene Expression in Trans-Generationally Immune Primed Bumblebee Workers Mimics a Primary Immune Response, PLoS One. 11 (2016) e0159635. https://doi.org/10.1371/journal.pone.0159635.

[141] A. Vilcinskas, The role of epigenetics in host–parasite coevolution: lessons from the model host insects *Galleria mellonella* and *Tribolium castaneum*, Zoology. 119 (2016) 273–280. https://doi.org/10.1016/j.zool.2016.05.004.

[142] P. Norouzitallab, K. Baruah, P. Biswas, D. Vanrompay, P. Bossier, Probing the phenomenon of trained immunity in invertebrates during a transgenerational study, using brine shrimp Artemia as a model system, Sci. Rep. 6 (2016) 21166. https://doi.org/10.1038/srep21166.

[143] J. Jussila, J. Makkonen, A. Vainikka, R. Kortet, H. Kokko, Crayfish plague dilemma: How to be a courteous killer?, Boreal Environ. Res. 19 (2014) 235–244.

[144] J.N. Thompson, Coevolution, Encycl. Life Sci. London, Nat. Publ. Gr. (2001).

[145] R.A. Schwenke, B.P. Lazzaro, M.F. Wolfner, Reproduction – Immunity Trade-Offs in Insects, (2017) 239–256. https://doi.org/10.1146/annurev-ento-010715-023924.Reproduction.

[146] L.L. Boštjančić, L. Bonassin, L. Anušić, L. Lovrenčić, V. Besendorfer, I. Maguire, F. Grandjean, C.M. Austin, C. Greve, A. Ben Hamadou, J. Mlinarec, The *Pontastacus leptodactylus* (Astacidae) Repeatome Provides Insight Into Genome Evolution and Reveals Remarkable Diversity of Satellite DNA, Front. Genet. 11 (2021). https://doi.org/10.3389/fgene.2020.611745.

[147] M.H. Tan, H.M. Gan, Y.P. Lee, F. Grandjean, L.J. Croft, C.M. Austin, A Giant Genome for a Giant Crayfish (*Cherax quadricarinatus*) With Insights Into cox1 Pseudogenes in Decapod Genomes, Front. Genet. 11 (2020). https://doi.org/10.3389/fgene.2020.00201.

[148] K. Koiwai, T. Koyama, H. Suzuki, R. Kawano, S. Tsuda, A. Toyoda, K. Kikuchi, L. Science, Single-cell RNA-seq analysis reveals penaeid shrimp hemocyte subpopulations and cell differentiation process, (2021) 1–27. https://doi.org/10.1101/2021.01.10.426076.

