## Supplementary figures and images for "Comparative transcriptome analysis of noble crayfish and marbled crayfish immune response to *Aphanomyces astaci* challenges"

### Figure S2.tif

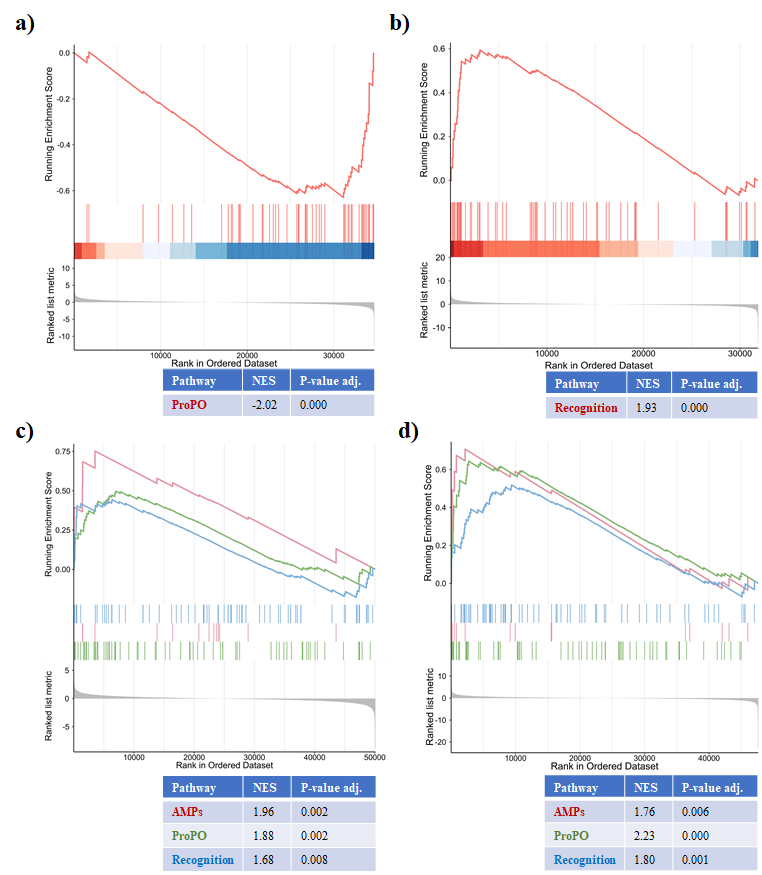
